# Functional and LC-MS/MS analysis of *in vitro* transcribed mRNAs carrying phosphorothioate or boranophosphate moieties reveal polyA tail modifications that prevent deadenylation without compromising protein expression

**DOI:** 10.1101/2020.07.02.184598

**Authors:** Dominika Strzelecka, Miroslaw Smietanski, Pawel J. Sikorski, Marcin Warminski, Joanna Kowalska, Jacek Jemielity

## Abstract

Chemical modifications enable preparation of mRNAs with augmented stability and translational activity. In this study, we explored how chemical modifications of 5’,3’-phosphodiester bonds in the mRNA body and polyA tail influence the biological properties of eukaryotic mRNA. To obtain modified and unmodified *in vitro* transcribed mRNAs, we used ATP and ATP analogues modified at the α-phosphate (containing either O-to-S or O-to-BH_3_ substitutions) and three different RNA polymerases—SP6, T7 and polyA polymerase. To verify the efficiency of incorporation of ATP analogues in the presence of ATP, we developed a liquid chromatography–tandem mass spectrometry (LC-MS/MS) method for quantitative assessment of modification frequency based on exhaustive degradation of the transcripts to 5’-mononucleotides. The method also estimated the average polyA tail lengths, thereby providing a versatile tool for establishing a structure-biological property relationship for mRNA. We found that mRNAs containing phosphorothioate groups within the polyA tail were substantially less susceptible to degradation by 3’-deadenylase than unmodified mRNA and were efficiently expressed in cultured cells, which makes them useful research tools and potential candidates for future development of mRNA-based therapeutics.

## 1. Introduction

In recent years, *in vitro* transcribed (IVT) mRNA has emerged as a promising candidate for therapeutic gene delivery.(1,2) Since proper mRNA function is dependent on its structural regulatory elements: a 5’ cap, 5’ and 3’ untranslated regions (UTRs), and the polyA tail at the 3’ end, rational modifications of these elements improve the potency of mRNA therapeutics. The use of both natural and unnatural chemical modifications of mRNA have paved the way to modulating stability, translational properties, and immunogenicity of IVT mRNAs (2–5). One of the structural elements that affect mRNA half-life and translation is the 5’ terminal 7-methylguanosine cap (6). Modifications of the 5’ cap may lead to augmented mRNA stability and expression in living cells (7–11). mRNA body modification via replacing uridine with pseudouridine or 1-methylpseudouridine has been found to enhance mRNA stability and translational properties, while decreasing immunogenicity (12–14). Introduction of particular sequences, such as the 3’ UTR of β-globin downstream of the open reading frame (ORF) provides similar effects (15–18). Improvement of translational properties has also been observed for mRNAs fluorescently labeled at the 3’ end of the polyA tail (19).

The polyA tail is the stretch of adenylate residues at the 3’ end of most of the mature mammalian mRNAs, (except histone-encoding replication-dependent genes (20)). PolyA tails of mature mammalian mRNAs consist of several dozens to up to 250 adenosine residues (21,22) and form cytoplasmic ribonucleoprotein (RNP) complexes with polyA-binding proteins (PABPs) (23). The interaction is necessary for efficient translation and controls mRNA stability (24–26). The polyA-PABP RNP complex interacts with the eIF4G scaffold protein stimulating the translation initiation process (27); thus, the presence of a polyA tail is essential for efficient protein expression. PolyA tails undergo gradual shortening (deadenylation) in an mRNA-specific manner (28), which is a necessary step before committing transcripts to one of the bulk mRNA decay pathways (29,30). Deadenylation is catalyzed by two major cytoplasmic deadenylase complexes: Pan2-Pan3, which initiates this process, and Ccr4-Not, acting at a later stage, finishing deadenylation prior to mRNA body decay (28,31). Structural features of polyA tail recognition by the Pan2-Pan3 deadenylase have been described, and the role of PABPs, together with polyA, in forming a specific binding scaffold for the Pan2-Pan3 heterotrimer has been revealed (32). Other studies revealed the possibility of stabilizing mRNA molecules by preventing deadenylation, which has been achieved by modifying the very 3’ end of mRNA (33,34). Herein, we envisaged that the chemical space for modifying mRNA and mRNA polyA tails was potentially much wider. We have previously found that modifying the triphosphate chain of the mRNA 5’ cap, e.g. by replacing one of the phosphate moieties with phosphorothioate or boranophosphate, has beneficial effects on mRNA stability and translation. However, modifications of phosphodiester bonds in the mRNA body or polyA tails have never been explored in this context. Here, we systematically investigated the effects of phosphodiester bonds modification on mRNA stability and translational properties to map out a detailed structure-activity relationship (SAR). To obtain phosphodiester-modified mRNAs (P-mod mRNAs), we utilized ATP analogs with non-bridging oxygen atom substitutions at the α-phosphate, i.e. ATPαS and ATPαBH_3_, which are enzymatically incorporated into RNA by means of RNA polymerases (35,36). However, this goal posed significant analytical challenges associated with the control and determination of polyA tail length and modification frequency. Thus, the development of proper analytical methods is a necessary prerequisite for further exploration of unnatural polyA tail modifications and precise determination of SAR in a manner previously achieved for the 5’ cap.

Several methods to measure polyA tail lengths have been reported, including the polymerase-based poly(A) length assay engaging capillary electrophoresis (37), ligation-mediated poly(A) test (LM-PAT), and RNase H-based assay (38). However, such methods are suitable for polyA tails of limited length due to poor resolution of the applied analytical methods, which is additionally complicated by polyA tail heterogeneity (39). The highest resolution sequencing method, such as TAIL-seq (40), PAlso-seq (41,42), and FLAM-seq (43) provide information on nucleotide sequence, polyA tail length, and the presence of adenine modifications. Mass spectrometry has been successfully used in the analysis of RNA, including mRNA, either by direct infusion analysis or combined with chromatographic techniques (44–51). This powerful technique provides information on RNA structure and composition, including polyA tail length. Although, it is a high resolution and sensitive method, the analysis of long mRNA may be difficult because of multiply charged states and associated spectra complexity (52).

In this study, to enable the analysis of P-mod mRNAs, we developed a new liquid chromatography–tandem mass spectrometry (LC-MS/MS) method that enabled simultaneous quantification of the number of modified phosphodiester bonds and estimation of the average polyA tail lengths in mRNA (Figure 1). Subsequently, we applied this method to characterize IVT P-mod RNAs carrying either phosphorothioate or boranophosphate modifications synthesized by three different RNA polymerases. Finally, LC-MS/MS characterized P-mod mRNAs were studied *in vitro* to evaluate their susceptibility to enzymatic deadenylation and in cultured cells to estimate the influence of phosphodiester modifications on protein expression. We obtained the first glimpse of the SAR of P-mod RNAs and identified modification patterns that were compatible with efficient protein expression.

**Figure 1.**
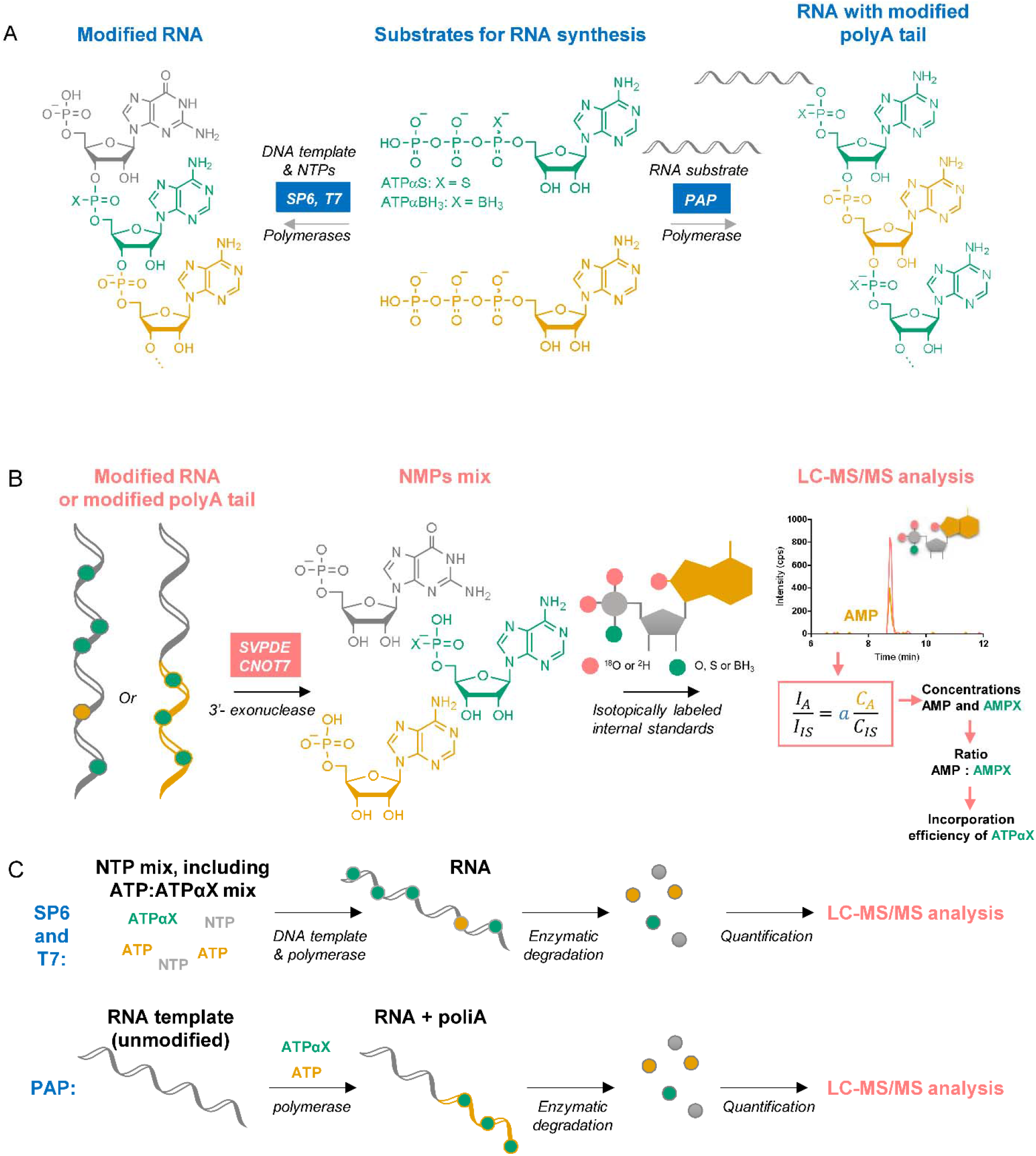
Synthesis and analysis of phosphate-modified RNA – experimental overview. (A) RNAs modified throughout the sequence are synthesized by *in vitro* transcription with the SP6 or T7 polymerase from a template also encoding the polyA tail in the presence of nucleoside triphosphates (NTPs) (left), whereas RNAs modified only within the polyA tail are synthesized by polyadenylation of unmodified RNA substrate with poly(A) polymerase (PAP) (right), (B) modified RNAs are subjected to exhaustive degradation by a mix of nucleases and the resulting mononucleotides (adenosine monophosphate (AMP), modified AMP, and guanosine monophosphate (GMP) as a reference) are quantified using liquid chromatography– tandem mass spectrometry (LC-MS/MS), (C) simplified workflow from nucleotide incorporation to RNA degradation and analysis.

## 2. Results

### 2.1. LC-MS/MS enabled determination of the phosphorothioate moiety content in RNA and estimation of an average polyA tail length

To enable systematic investigation of the SAR of modified RNAs, we developed an LC-MS/MS method for quantitative assessment of the incorporation of phosphate-modified ATP analogs into transcripts by RNA polymerases (Figure 1). If ATPαS was utilized by the polymerase as a substrate, a modified RNA (modRNA) with phosphorothioate bonds was produced (Figure S4). For polymerases that have been studied so far, only the *S*_P_ isomer of ATPαS has been accepted as a substrate, and the reaction proceeds with inversion of the configuration at the phosphorus atom (65). To assess the efficiency of ATPαS incorporation, modRNA was enzymatically degraded to single NMPs and adenosine 5’-phosphorothioate (AMPS). Ion pair chromatography coupled with ESI and a triple quadrupole analyzer was then used to resolve and quantify AMPS and select NMPs. Synthetic isotopologs of RNA degradation products labeled with heavy oxygen (^18^O) within the phosphate were used as internal standards to facilitate quantification. The data were then mathematically processed to determine the absolute concentrations of these nucleotides, the AMP/AMPS ratio, and, if a RNA of known sequence with a 3’ terminal polyA tract was analyzed, the average polyA tail length was also estimated. We found that the key factor during method development was the optimization of the RNA degradation procedure to ensure complete RNA degradation to (modified) monophosphates, without further decomposition of released nucleotides.

To obtain isotopically labeled internal standards, chemical (thio)phosphorylation of nucleosides followed by hydrolysis with heavy water was applied (Figure S1) (57). Using this method, the internal standards containing two heavy oxygen (^18^O) atoms in the phosphate group were synthesized for all tested NMPs and AMPS. Heavy AMPS, AMP and GMP were used as internal standards for quantification of, respectively, AMPS, AMP, and GMP released during RNA degradation (Figure 2A). Since guanine was present only in the main RNA body (and not the polyA tail), the quantity of GMP was used to normalize the amounts of RNAs that were analyzed, which was particularly important when comparing samples with different (or unknown) polyA tail lengths. The detection of AMPS and other NMPs was performed in MRM mode. MRM pairs were selected for each analyte and an internal standard based on the determined fragmentation pattern (Figure 2B, Table S1). The predominant fragmentation reaction for AMPS was the 362 → 95 transition, corresponding to the cleavage of the 5’-phosphorothioate bond and release of the PSO_2_^-^ ion (57). The chromatographic conditions were also optimized to achieve good separation and a satisfactory peak shape for all studied analytes. To that end, ion pair chromatography with N’,N’-dimethylhexylamine as a mobile phase reagent (56) has provided the best results in our hands (Figure 2C). Calibration curves were subsequently determined for each analyte under optimized conditions (Figure 2D, Figure S2). Based on the calibration curves, limits of quantification (LOQ) were determined (Table S3). The final step of method development was establishing RNA degradation conditions. The goal was to achieve complete RNA degradation within a reasonable time, regardless of the modification content. This task was not trivial, since it has been known that the phosphorothioate modification, in general, stabilizes RNA against nucleolytic cleavage(66,67). Moreover, we found that high concentrations of nucleolytic enzymes in the RNA sample leads to partial desulfurization of AMPS to AMP. Therefore, we tested different concentrations and mixtures of nucleolytic enzymes. We found that the optimal results were achieved for the mix of phosphodiesterase I *Crotalus adamanteus* venom (SVPDE) and human CNOT7 deadenylase. SVPDE, which is a commercially available 3’→5’ exonuclease with broad spectrum specificity, releases 5’-mononucleotides (68); the enzyme is capable of degrading modified RNA (69). The concentration of SVPDE in our assay was initially optimized using unmodified IVT RNA A and a fully modified RNA A. As expected, a higher enzyme concentration was required to completely degrade modified RNA A. However, during LC-MS/MS analysis of nucleotides released from fully modified RNA A, AMP was also found in the sample (Figure 2E). This suggested that SVPDE at high concentrations also catalyzes AMPS desulfurization, which has been previously reported in the literature.(65) Hence, we additionally optimized degradation by adjusting enzyme concentrations to maximize the AMPS to AMP ratio and maintain complete degradation of RNA (Figure 2F,G). Next, we applied similar conditions to the degradation of longer RNAs, including mRNAs (RNA F2). We found that polyadenylated mRNAs were less susceptible to SVPDE compared to other RNAs. Therefore, to ensure complete degradation of polyadenylated RNAs, we used a mixture of SVPDE and CNOT7, which has 3→5’ polyA exoribonuclease activity (70). These conditions enabled complete degradation of most of the analyzed polyadenylated RNAs, while maintaining low desulfurization levels (Figure S3). To verify the extent of AMPS desulfurization under optimized digestion conditions, RNA E, which was devoid of an A in the sequence, was polyadenylated with a mixture of deuterated ATP (^D^ATP) and ATPαS D1, following RNA degradation and LC-MS analysis. The amount of undeuterated AMP, which could be derived only from AMPS was approximately 10% (Figure S5). For each kind of RNA, the completeness of the digestion process was additionally verified by gel electrophoresis. If any undigested RNA was observed, the digestion conditions were adjusted individually, which was usually necessary for fully modified RNA.

**Figure 2.**
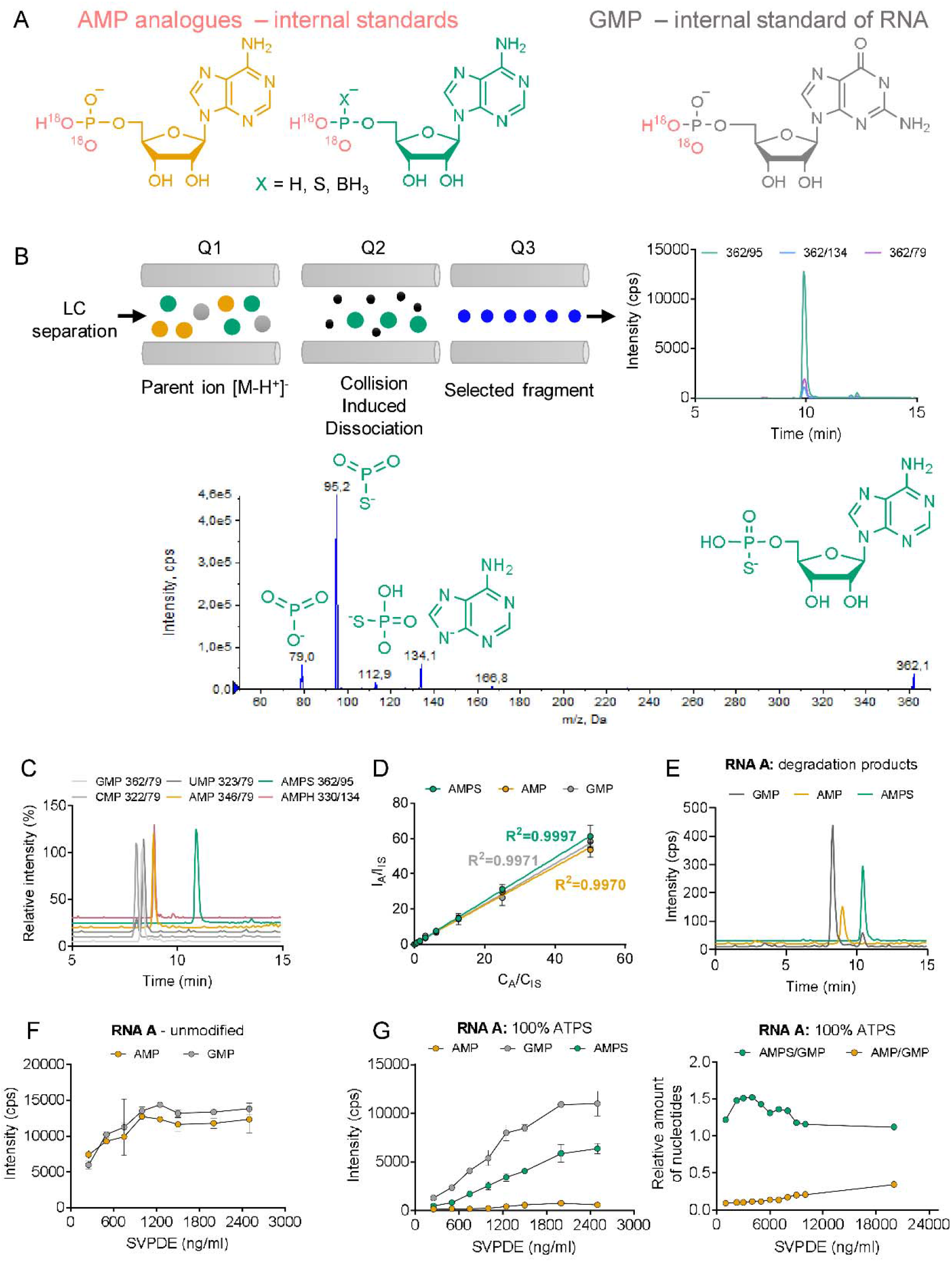
Liquid chromatography–tandem mass spectrometry (LC-MS/MS) method development. (A) Synthesis of isotopically labeled internal standards (heavy adenosine monophosphate (AMP), heavy adenosine 5’-O-monothiophosphate (AMPS), heavy guanosine monophosphate (GMP), heavy adenosine-5’-O-(H-phosphonate) (AMPH); (B) identification and optimization of suitable multiple reaction monitoring (MRM) pairs; (C) optimization of LC separation conditions; (D) determination of calibration curves; (E) high performance liquid chromatography (HPLC)-MS analysis of an authentic RNA sample (phosphorothioate RNA A); (F,G) optimization of nuclease (snake venom phosphodiesterase (SVPDE)) concentrations to ensure complete degradation of RNA A (unmodified, F or modified with phosphorothioate moieties, (G).

For polyadenylated RNAs, the average polyA tail length was also estimated (Figure S6, 5-8). To that end, the ratio of AMP and GMP concentrations (*r*_ref_) in the non-polyadenylated RNA sequence was experimentally determined. Then, the average polyA tail length was calculated based on Equation 3. To initially verify the reliability of the method, we tested a chemically synthesized short RNA fragment (RNA C) containing a single G moiety and a string of adenosines (Figure S3A). The determined AMP to GMP ratio was comparable with the value expected based on the sequence.

### 2.2. *S*_p_-ATPαS was a substrate for T7 and SP6 polymerases

We next tested how efficiently the ATP analogs were incorporated into RNA during *in vitro* transcription and polyadenylation compared to unmodified ATP. Based on previously reported knowledge, we have assumed that *S*p-ATPαS (isomer D1) will be incorporated into RNA, while *R*_P_-ATPαS (isomer D2) will be neither a substrate, nor an inhibitor (71). The *in vitro* transcription reactions were performed from a DNA template containing the promoter followed by a sequence of 35 nt using a standard protocol for SP6 RNA polymerase or T7 RNA polymerase. The reactions contained different ratios of ATP to ATPαS (either D1 or D2 isomer). The yield and integrity of transcription products were analyzed by gel electrophoresis (Figure S4). High yields of IVT products were obtained in the presence of ATPαS D1 independently of the ATP:ATPαS D1 ratio used in the reaction with SP6 polymerase. In contrast, the presence of higher concentrations of ATPαS D2 isomer resulted in a lower transcription yield. The RNAs were then subjected to degradation by SVPDE and subjected to LC-MS/MS analysis (Figure 3). The determined concentrations of AMP, AMPS and GMP were plotted as a function of ATPαS D1 or D2 used in the *in vitro* transcription reaction (Figure 3A,B) and the data were normalized to GMP (Figure 3C,D). The AMPS to AMP ratio corresponded to the ATPαS D1 to ATP ratio (Figure 3A), indicating that ATPαS D1 incorporation efficiency was similar to that of ATP (Figure 3E). In contrast, regardless the ATPαS D2:ATP ratio, AMPS was barely detected in RNA degradation products (Figure 3B). These findings are in good agreement with previous reports on the stereoselectivity of T7 RNA polymerase, which revealed that only isomer D1 is incorporated (35,71). Next, we compared SP6 and T7 polymerase activities. To that end, the corresponding enzymatic reactions were performed by SP6 and T7 RNA polymerases in the presence of a 1:1 ATP:ATPαS ratio (Figure 3F). We found that under the same conditions, T7 polymerase incorporated ATPαS D1 at a slightly lower frequency than SP6, whereas ATPαS D2 was not incorporated.

**Figure 3.**
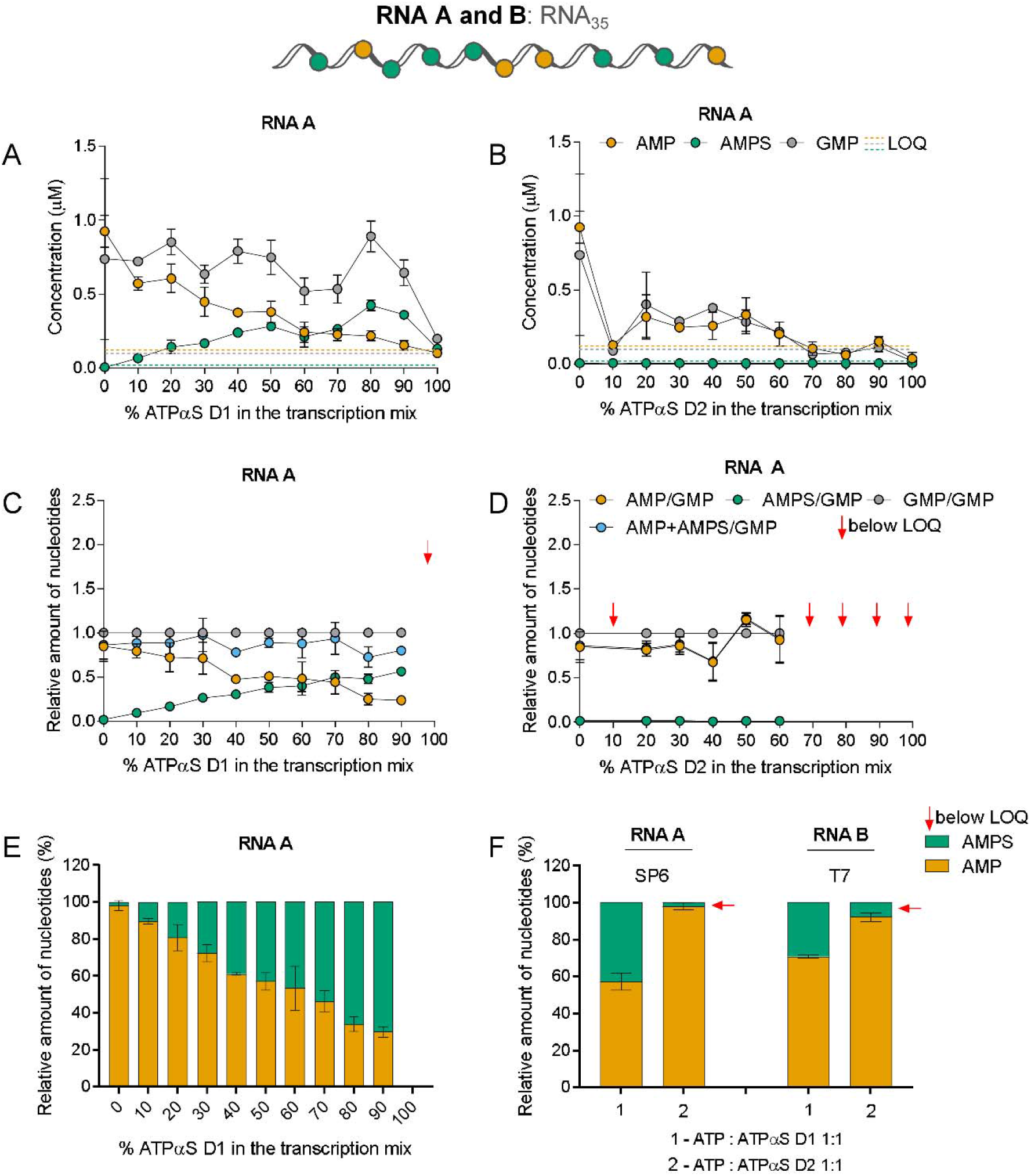
SP6 and T7 polymerases accept ATPαS D1 as a substrate. Short RNAs (RNA A or B) were obtained by *in vitro* transcription by SP6 or T7 RNA polymerase, respectively, in the presence of different ATP:ATPαS D1/D2 ratios. *In vitro* transcribed (IVT) RNAs (20 ng) were incubated with 3 μg/mL snake venom phosphodiesterase (SVPDE) for 1 h at 37 °C (see Materials and Methods for further details). Concentrations of nucleotides after RNA digestion are determined based on liquid chromatography-tandem mass spectrometry (LC-MS/MS) analysis (A, B). The data are normalized to guanosine monophosphate (GMP) to account for differences in transcript amounts that were analyzed (C, D). The phosphorothioate moiety frequency in adenine nucleotides as a function of ATPαS D1 concentration in the transcription mix is shown in (E), (F) shows the frequency of phosphorothioate moieties in adenine nucleotides found in RNAs obtained with SP6 and T7, and ATPαS (either D1 or D2). Data points represent mean values from triplicate experiments ± standard deviation (SD).

### 2.3. High concentrations of ATPαS D1 in the transcription mix reduced the transcription yield and did not augment mRNA translational properties

After establishing that both SP6 and T7 RNA polymerases incorporated ATPαS D1 into RNA, we determined how this phosphorothioate modification influenced the biological properties of mRNAs. This issue has rarely been investigated in the literature, and to the best of our knowledge, only in the context of prokaryotic mRNA (72).

To this end, modified mRNAs encoding *Firefly* luciferase as a reporter gene (RNA F) were prepared by *in vitro* transcription. The template for the *in vitro* transcription reaction encoded a short 5’ UTR sequence, followed by the *Firefly* luciferase ORF, two consecutive *H. sapiens* β-globin 3’ UTRs, and a 128 nt long (A_128_) polyA tail (Figure 4A). Considering the determined stereoselectivity of T7 and SP6 RNA polymerases (Figure 3), we used the ATPαS D1 stereoisomer as a substrate for RNA modification. The typical *in vitro* transcription reaction contained a mix of unmodified NTPs (0.5 mM CTP, UTP, 0.125 mM GTP, and 1.25 mM cap analog – β-S-ARCA), and a mix of ATP and ATPαS D1, at a total concentration of 0.5 mM and varying molar ratios (10:0, 9:1,2:1, 1:1, 1:4, or 0:10). To compare the transcription yields and quality of products, the transcripts were purified on a silica-membrane and analyzed on an agarose gel (Figure 4B). The transcription yield decreased with an increasing concentration of ATPαS D1 in the *in vitro* transcription mix. This suggested that ATPαS D1 was either a less efficient substrate of T7 and SP6 RNA polymerases or inhibited the activity of these RNA polymerases at higher concentrations.

**Figure 4.**
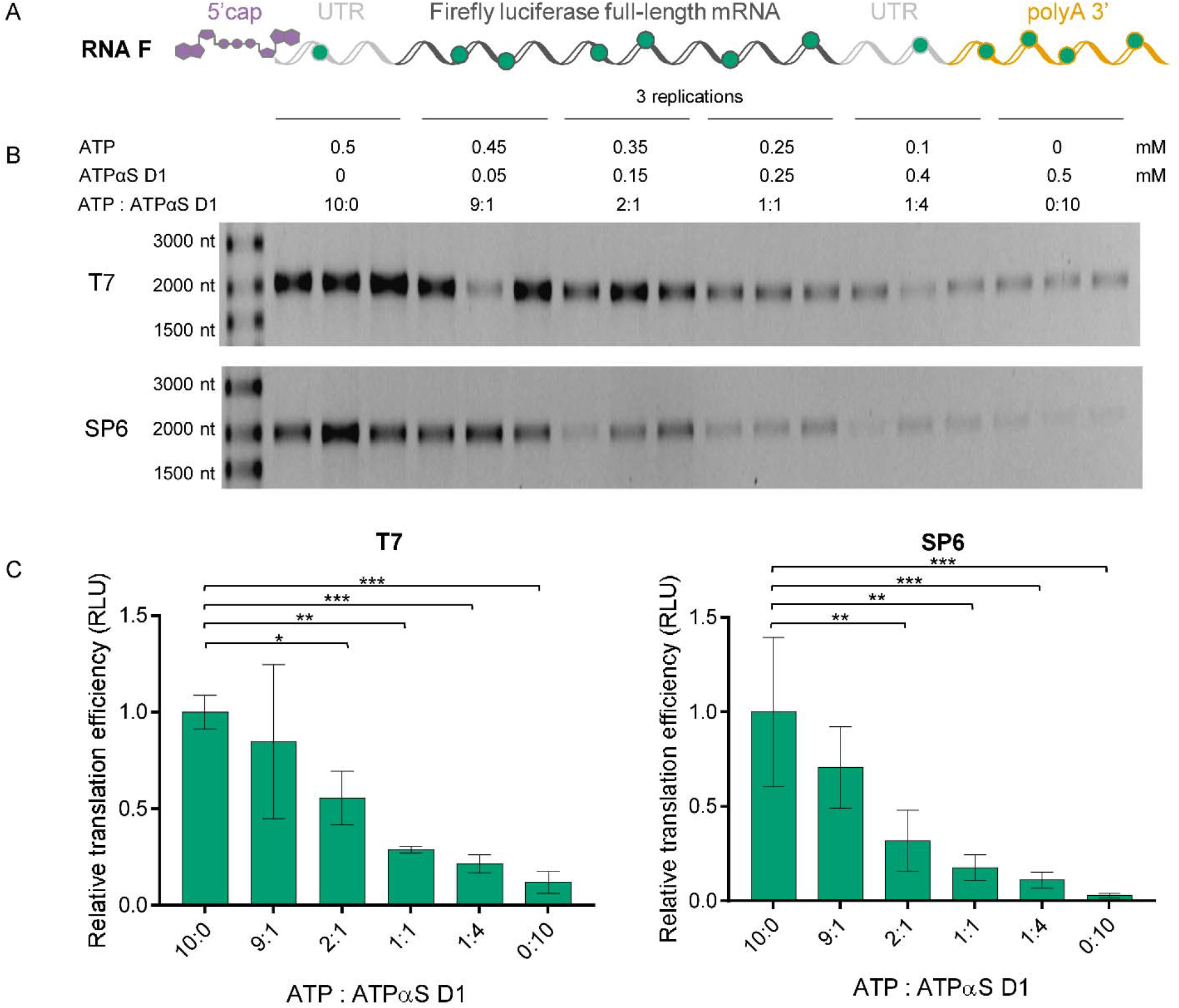
Analysis of homogeneity and translational properties of mRNAs uniformly modified with phosphorothioate moieties. (A) Schematic representation of *Firefly* luciferase (RNA F) *in vitro* transcribed (IVT) with T7 or SP6 RNA polymerase. Green dots represent phosphorothioate moieties randomly placed within the mRNA body; (B) RNA F variants co-transcriptionally 5’-capped with the cap analog (β-S-ARCA D1) and obtained in the presence of the indicated ATP:ATPαS D1 molar ratios (10:0, 9:1,2:1, 1:1, 1:4, or 0:10) are synthesized with T7 or SP6 RNA polymerase and purified on a silica membrane (NucleoSpin RNA Clean-up XS, Macherey-Nagel). The transcription yields and quality of transcripts are analyzed by electrophoresis on a 1% agarose gel. (C) Translation efficiencies are tested in rabbit reticulocyte lysate (Promega) using appropriate RNA F concentrations (3, 1.5, 0.75, and 0.375 ng/μL) (see Materials and Methods for further details). Translation efficiencies are determined based on measured luminescence as a function of RNA F concentration. Slopes determined for different variants of RNA F are normalized to the slope of unmodified RNA F (10:0), calculated as the mean value from three independent experiments ± standard deviation (SD). Statistical significance: * *p* < 0.05, ** *p* < 0.01, *** *p* < 0.001, **** *p* < 0.0001 (one-way analysis of variance (ANOVA) with Dunnett’s multiple comparison test). Only statistically significant differences are marked in the graph.

RNA variants with different amounts of phosphorothioate modifications were then evaluated for their translation efficiency in the rabbit reticulocyte lysate. The lysate, optimized for cap dependent translation, was programmed with RNA F and the amount of luciferase was determined after 60 min by luminometry. The translation efficiencies of RNA F decreased with an increasing concentration of ATPαS D1 used in the transcription mix (Figure 4C). RNAs obtained in the presence of solely ATPαS D1 (i.e. in the absence of ATP) were translated at 10-fold lower efficiency than unmodified transcripts.

Overall, these results indicated that randomly modifying the entire mRNA sequence with phosphorothioate moieties reduced both the transcription yield and translational activity of mRNA. We envisaged that the most likely interferences with the translation processes resulted from the modifications within the ORF or UTRs, which may impair ribosomal movement. Thus, in the next steps of the study, we investigated properties of mRNAs modified solely within the polyA tails.

### 2.4. PAP catalyzed the synthesis of phosphorothioate-modified polyA tails

To achieve selective polyA tail modification of IVT RNAs, we used PAP from *E. coli,* a commercially available enzyme that specifically recognized the 3’ end of single-stranded RNA (ssRNA) and performed template-independent synthesis of single-stranded polyA tracts using ATP as a substrate. Since it had not been previously investigated whether PAP accepted phosphate-modified ATP analogs as substrates, we first tested its activity towards ATPαS D1 and D2. In a pilot experiment, polyadenylation reactions of RNA A (35 nt) were performed. To this end, three PAP concentrations (1U, 2U, 4U) and either ATP alone, ATP:ATPαS in a 1:1 ratio, or ATPαS alone were tested (Figure 5). The obtained products were initially analyzed by agarose gel electrophoresis, which enabled visual estimation of the reaction efficiency and the dominant polyA tail length (Figure 5A,B). In the presence of ATP alone, polyadenylated RNAs of similar visual quality and dominant lengths between 300–500 nucleotides were formed in the presence of all tested PAP concentrations. In the presence of 1:1 ATP:ATPαS D1, the elongation of RNA was also observed, but the resulting RNAs appeared to be shorter and more homogenous, which indicated either a shorter length, disturbed migration due to modifications, or a smaller length distribution. In the presence of 100% ATPαS D1, the formed polyA tails migrated even faster (at the level corresponding to unmodified RNA). In the presence of the ATP:ATPαS D2 mix, the polyadenylation of RNA was clearly visible, and the resulting transcripts migrated only slightly faster than those obtained in the presence of ATP. In contrast, in the presence of 100% ATPαS D2, RNA elongation was not observed. Independently, the obtained transcripts were digested using a SVPDE and CNOT7 mixture and the obtained products were analyzed by LC-MS/MS to determine RNA composition, polyA composition, and to estimate the average polyA tail length. The amounts of AMP and AMPS released from RNA were initially normalized to the GMP amount (Figure 5C,D). Knowing the concentrations and relative ratios of AMPS, AMP and GMP in the samples, as well as the template sequence (specifically number of As and Gs in the template), the average polyA tail length was estimated using Equations 3 and 4 (Material and Methods; Figure 5E,F), which also enabled estimation of the polyA tail composition (Figure 5G,H). AMPS was detected as a product of RNA degradation if ATPαS D1 was present in the polyadenylation reaction. RNAs obtained in the presence of ATP:ATPαS D1 released significant amounts of AMPS, corresponding to equally efficient incorporation of ATP and ATPαS D1 (Figure 5C). The average lengths of polyAs in those RNAs (45–95 nt) were comparable or only slightly shorter than those in corresponding RNAs obtained in the presence of ATP, and increased with increasing amounts of PAP (Figure 5E). In contrast, polyA tails synthesized in the presence of 100% ATPαS D1 were notably shorter (~5 nt). The quantification of products resulting from degradation of these indicated a notable amount (~40%) of AMP present in the polyA (Figure 5G); however, this was likely due to a method limitation related to very short polyA tails. Overall, we concluded that ATPαS D1, was accepted as a PAP substrate, but the incorporation process was efficient only if ATP was also present in the reaction mixture. If RNA was polyadenylated in the presence of the ATP:ATPαS D2 mix, AMP was detected among the degradation products, while AMPS was not observed (Figure 5D). The average RNA lengths were comparable to those obtained in the presence of ATP (Figure 5F). RNAs incubated with PAP in the presence of 100% ATPαS D2 did not contain polyA tails (Figure 5F). This suggested that ATPαS D2 was not accepted as a substrate for PAP, similar to that for the T7 and SP6 polymerases.

**Figure 5.**
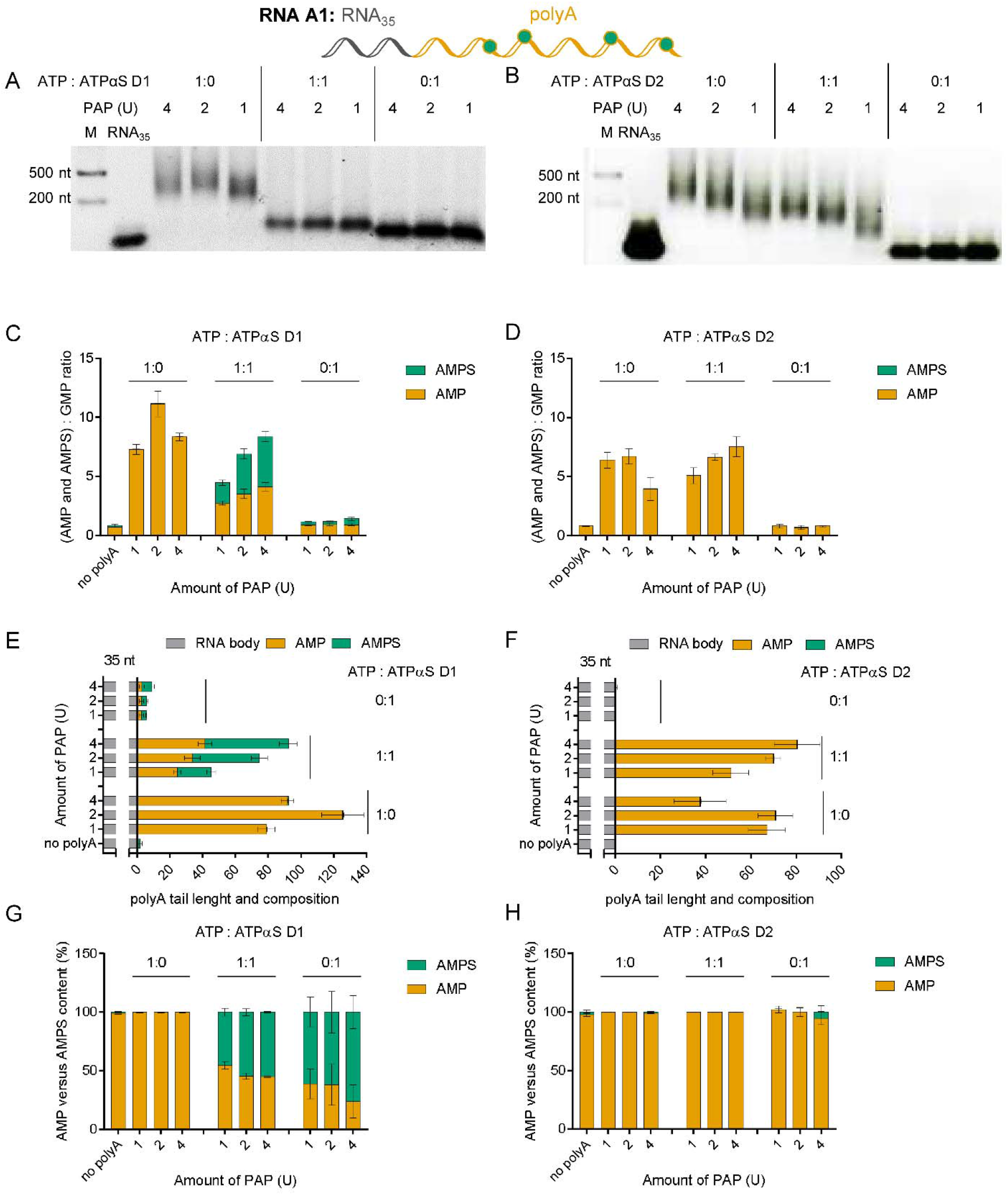
PolyA polymerase (PAP) incorporates ATPαS D1 in the presence of ATP to produce modified polyA tails. Analysis of short RNAs with polyA (RNA A1) tails added by PAP using different ATP:ATPαS ratios (1 mM). Short RNAs (RNA A) are obtained by SP6 polymerase. RNA A1 (20 ng) is digested using the snake venom phosphodiesterase (SVPDE) and Ccr4-Not transcription complex subunit 7 (CNOT7) enzymes for 1 h at 37 ° C (see Materials and Methods for further details). Data points represent mean values from triplicate experiments ± standard error of the mean (SEM). (A, C, E, G) Data obtained for ATPαS D1, (B, D, F, H) data obtained for ATPαS D2. Data shown are from a single agarose gel (A, B). RNA composition is determined based on liquid chromatography–tandem mass spectrometry (LC-MS/MS) quantification analysis followed by normalization to guanosine monophosphate (GMP) (C, D). PolyA tail length and composition (E, F), as calculated by the procedure described in Materials and Methods. Incorporation efficiency of ATP versus ATPαS D1 or D2 by PAP is calculated (G, H).

The experiments also revealed a notable inconsistency in the lengths of polyA tails estimated by LC-MS/MS and PAGE analysis for RNAs obtained in the presence of ATP. To verify whether this arose from significant polyA tail length heterogeneity or an inaccuracy in our MS/MS method, we prepared a set of unmodified mRNAs (RNA F2) carrying polyA tails of various lengths (different polyA tail lengths were achieved by varying the duration of the polyadenylation reaction) and analyzed them using both methods (Figure S6). We found that RNA lengths determined by LC-MS/MS were around 3-times shorter than the RNA length estimated by electrophoretic analysis, but the correlation between the methods remained linear. Hence, we hypothesized that these differences between the methods arose from large RNA heterogeneity, which caused a significant difference between dominant and average mRNA lengths (Figure S6H).

### 2.5. Phosphorothioate modifications decreased polyA tail susceptibility to deadenylation and slightly influenced protein expression levels in cells

To study the effects of polyA tail modification on mRNA stability and translation, we prepared unmodified mRNAs (RNA G and RNA H) and enzymatically added a polyA tail onto the 3’ end using PAP and phosphate-modified ATP analogs (Figure 6A, Figure 8A). We investigated various properties of mRNAs with modified polyA tails, including polyA tail length, composition, susceptibility to deadenylation, and protein expression efficiency in mammalian cells. The DNA templates for *in vitro* transcription used in these experiments encoded a short 5’ UTR sequence, followed by the *Gaussia* luciferase ORF and either a single or double *H. sapiens* β-globin 3’ UTR sequence (RNA H and RNA G, respectively), and in the case of the reference mRNA (RNA I), a A_128_ polyA tail was encoded. To avoid interference from doublestranded RNA (dsRNA) impurities, the IVT mRNAs have been purified by RP HPLC prior to polyadenylation (73,74). This procedure significantly increased the homogeneity of mRNAs used in the polyadenylation step, thereby facilitating LC-MS/MS analysis, since the presence of any additional RNAs in the PAP reaction mix affected the accurate concentration determination of total polyadenylated mRNA. RP HPLC-purified RNA G co-transcriptionally capped with ARCA (Figure S7A,B) was polyadenylated by PAP in the presence of either ATP or an ATP:ATPαS D1 mix (at 9:1, 4:1, 3:1, 2:1, and 1:1 molar ratios). The polyadenylation protocol was optimized (Figure S8) to yield RNA G1 with ~200–250 nucleotide long (based on agarose gel) polyA tails containing a phosphorothioate modification as the predominant product. During initial polyadenylation experiments, we noted that high concentrations of ATPαS D1 in the polyadenylation reaction decreased the homogeneity of RNA G1 (Figures S9, S10); therefore, the 1:1 molar ratio of ATP:ATPαS D1 was not exceeded.

**Figure 6.**
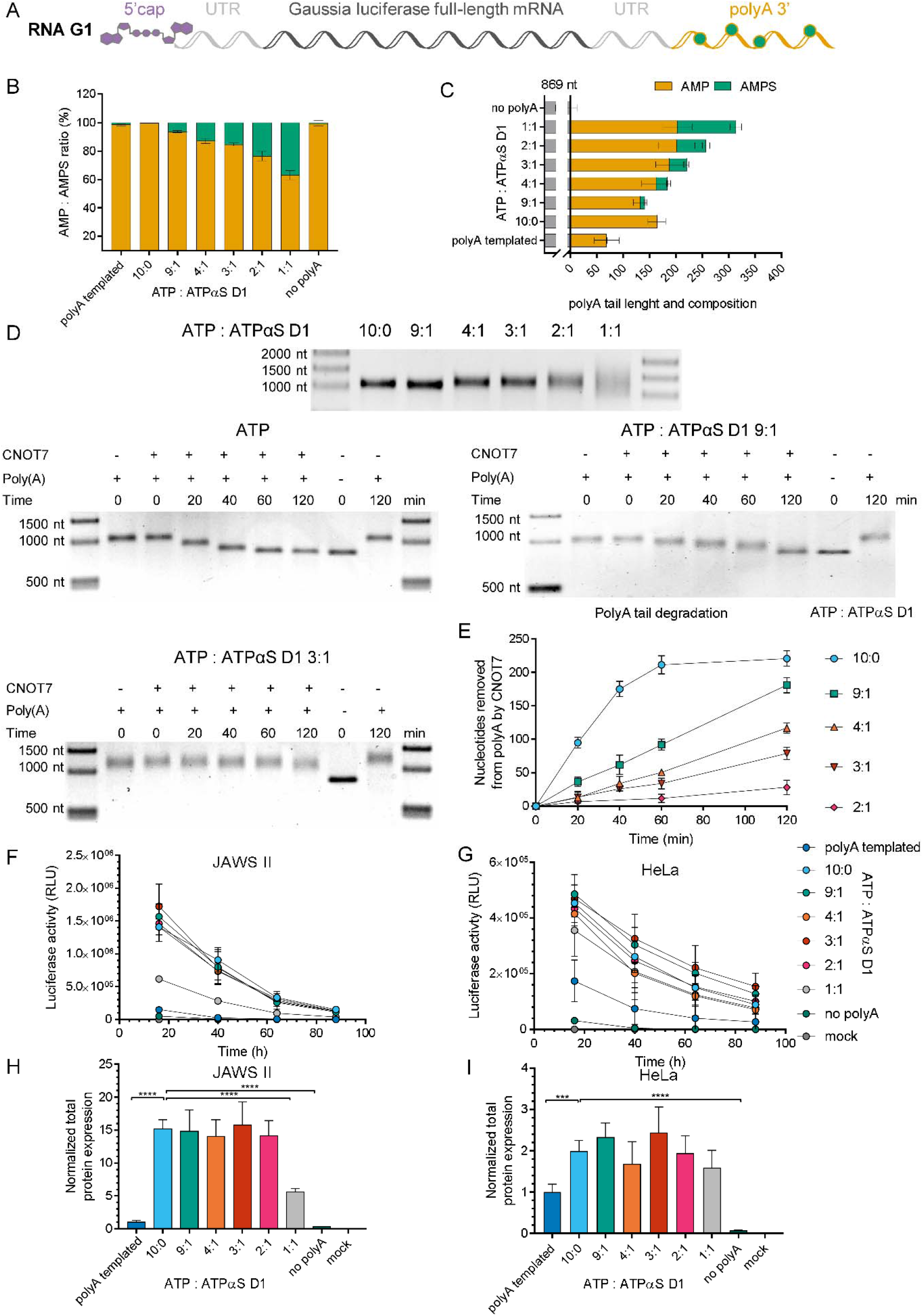
Analysis of mRNAs carrying phosphorothioate moieties within the polyA tail. (A) Schematic representation of the *Gaussia* luciferase-coding transcript, RP HPLC-purified and polyadenylated with polyA polymerase (PAP) in the presence of ATP or various mixtures of ATP:ATPαS D1 (9:1, 4:1, 3:1,2:1, or 1:1 molar ratio). (B) RNA G1 polyA tail composition (the mean ± standard error of the mean (SEM)), and (C) average polyA tail lengths (the mean ± SEM) obtained from liquid chromatography–tandem mass spectrometry (LC-MS/MS). (D) Analysis of RNA G1 susceptibility to deadenylation. RNA G1 variants are incubated with Ccr4-Not transcription complex subunit 7 (CNOT7) deadenylase for 0–120 min, and the polyA tail degradation rate is analyzed on a 1% agarose gel using Image Lab 6.0.1 Software (Bio-Rad). The length of the polyA tail at each time point is estimated as the difference between digested RNA G1 and RNA G (transcript before polyadenylation). (E) Susceptibility to deadenylation of RNA G1 polyadenylated with various ATP:ATPαS D1 ratios, estimated as the number of nucleosides removed by CNOT7 deadenylase at particular time points. (F-I) Translational properties of RNA G1 with phosphorothioate-modified polyA tails. *Gaussia* luciferase activity in the supernatant of (F) JAWS II and (G) HeLa cells, as measured 16, 40, 64, and 88 h after transfection with RNA G1. The cell medium was exchanged after each measurement. Data points represent mean values ± the standard deviation (SD) of one biological replicate consisting of three independent transfections. Total protein expression (cumulative luminescence) over 4 days in (H) JAWS II and (I) HeLa cells calculated from the same experiment. Bars represent the mean value ± SD normalized to RNA I (with a template-encoded A_128_ polyA tail). Statistical significance: * *p* < 0.05, ** *p* < 0.01, *** *p* < 0.001, **** *p* < 0.0001 (one-way analysis of variance (ANOVA) with Dunnett’s multiple comparison test). Only statistically significant differences are marked on the graph.

The RNAs were first analyzed by LC-MS/MS to estimate the polyA tail composition and average length. The results revealed that ATPαS D1 was efficiently incorporated into the polyA tail by PAP (Figure 6B, Table S4). The frequency of AMPS in the polyA tail, as determined using LC-MS/MS, was approximately 60–70% of the expected phosphorothioate moiety content, according to the molar ratio of ATP:ATPαS D1 used in the polyadenylation reaction (Table S4). The average polyA tail lengths determined by LC-MS/MS were similar to the lengths estimated by agarose gel electrophoresis (Figure 6C, Table S5). Moreover, the sensitivity of LC-MS/MS allowed for the analysis of the average polyA tail length of RNA G1, polyadenylated by PAP with a 1:1 molar ratio of ATP:ATPαS D1, whereas the length of the RNA G1 variant was difficult to estimate by electrophoretic analysis, due to heterogeneity (Figure 6D, Figures S9, S10).

To analyze the susceptibility to enzymatic degradation of modified polyA tails in mRNA, we set-up a deadenylation assay using CNOT7 deadenylase. CNOT7 is one of two catalytic subunits in the human Ccr4-Not deadenylase complex responsible for deadenylation of mRNAs in the cytoplasm (29). To this end, we performed time-course experiments to obtain the complete degradation of unmodified polyA tails, and to verify that recombinant CNOT7 was able to degrade only the polyA tail and leave the mRNA body intact (Figure S11). Next, we used this protocol for deadenylation time-course experiments using RNA G1 variants with polyA tails synthesized by PAP and selected ATP:ATPαS D1 molar ratios (9:1, 4:1, 3:1,2:1, and 1:1) (Figure 6D, Figures S9, S10). We found that RNA G1 variants with polyA tails containing phosphorothioate modification were significantly less susceptible to CNOT7 deadenylase than unmodified RNA G1 (Figure 6D,E; Table S6). RNA G1 with a polyA tail obtained at a 9:1 ATP:ATPαS D1 ratio, which contained 6% phosphorothioate moieties in the polyA tail (according to LC-MS/MS analysis), was degraded by CNOT7 at a rate three times slower than unmodified RNA G1 (V_0_ 1.5 nt/min and V_0_ 4.3 nt/min, respectively). Further increases in the phosphorothioate content in the polyA tail to 13%, 15%, and 23% corresponded to a deadenylation rate reduced by 4.4-fold, 6.7-fold and almost 20-fold, respectively (Table S4).

We next investigated the influence of phosphorothioate moieties present in the polyA on mRNA expression in living cells. To facilitate cell culture experiments, we used *Gaussia* luciferase as a reporter protein, since this type of luciferase is a secretory enzyme with a long half-life (~6 days), enabling straightforward monitoring of time-dependent mRNA expression by quantification of luminescence in cell culture medium (75). Thus, variants of RNA G1 were transfected into two mammalian cell lines: HeLa and JAWS II representing tumor and non-tumor dendritic cells (DCs), respectively. *Gaussia* luciferase activity was measured over a period of 4 days (Figure 6F,G). In both cell lines the total protein expression, reflected by luciferase activity, was similar for all tested mRNAs suggesting that phosphorothioate modification did not affect expression of the reporter protein (Figure 6H,I). Interestingly, both in HeLa and JAWS II cells, protein expression for unmodified RNA G1 was more efficient than for the unmodified RNA I used as a reference, which was consistent with the presence of a shorter polyA tail on RNA I (Figure 6H,I). It was also noted that similar protein expression levels for most of the RNA G1 variants with phosphorothioate-modified polyA tails did not correspond to decreased susceptibility to deadenylation, as determined with CNOT7 deadenylase.

These results altogether suggested that: (i) using a mixture of ATP and ATPαS D1 for polyadenylation provided access to mRNAs with phosphorothioate-modified polyA tails; (ii) phosphorothioate moieties decreased polyA tail susceptibility to deadenylation by CNOT7; and (iii) protein expression levels in JAWS II cells did not change with increasing phosphorothioate moiety content in the polyA tails of transfected mRNA.

### 2.6. Boranophosphate RNAs released AMPH as a degradation product

After demonstrating the utility of the method to analyze RNA with a phosphorothioate moiety, we determined whether it could be adapted to analyze RNAs carrying boranophosphate modifications. The boranophosphate moiety has similar structural properties and net charge to phosphorothioate, and also exhibits nuclease resistance (76). During the optimization of the method, we observed that the synthesized heavy AMPBH_3_, was chemically unstable (Figure S12A,B) and thus, could not be used as an internal standard. Furthermore, the main product of boranophosphate RNA degradation was not the expected AMPBH_3_, but adenosine *H*-phosphonate (AMPH) (Figure S12C). As such, heavy AMPH was used as an internal standard for quantification of both AMPH and AMPBH_3_ in the samples. AMPH has similar fragmentation pathways to AMPBH_3_ (57), but is chemically stable. Additionally, using heavy AMPH as an internal standard eliminated the problem of isotopic abundance of boron (20% ^10^B and 80% ^11^B), which decreased the sensitivity of detection.

To investigate the incorporation of boranophosphate analogs of ATP by PAP, we prepared RNAs that were depleted of adenosines throughout the sequence (RNA E). RNA E was obtained by *in vitro* transcription in the presence of T7 polymerase and subjected to a polyadenylation reaction, in which either 8-deutero ATP (^D^ATP) was used instead of ATP, or a mixture of ^D^ATP:ATPαBH_3_ (D1 or D2) was used. Polyadenylated RNA E1 was digested using a mixture of SVPDE and CNOT7, and the products were analyzed by LC-MS/MS (Figure 7). AMPH was observed as the main unnatural product released from RNA E1, obtained in the presence of ATPαBH_3_ D1 (1:1 ^D^ATP:ATPαBH_3_ D1 that was used for the polyadenylation reaction), while AMPBH_3_ was barely observed (Figure 7C). RNAs obtained in the presence of 100% ATPαBH_3_ D1 also released AMPH, but the quantified polyA tails were ~ 10-fold shorter compared to conditions including only ATP. Neither AMPH nor AMPBH_3_ were detected in samples generated from RNA E1 obtained using ^D^ATP and a ATPαBH_3_ D2 in a 1:1 ratio, which indicated that ATPαBH_3_ D2 was not incorporated by PAP. Compared to ATPαS D1, ATPαBH_3_ D1 seemed to be incorporated with lower frequency; ~40% AMPS (Figure S5) and ~20% AMPH (Figure 7) were detected when a 1:1 ratio of ATP and ATPαX isomer D1 was used for polyadenylation. Overall, the results indicated that only ATPαBH_3_ D1 was recognized as a substrate by PAP, but the polymerization reaction was less efficient than that for ATP and ATPαS D1.

**Figure 7.**
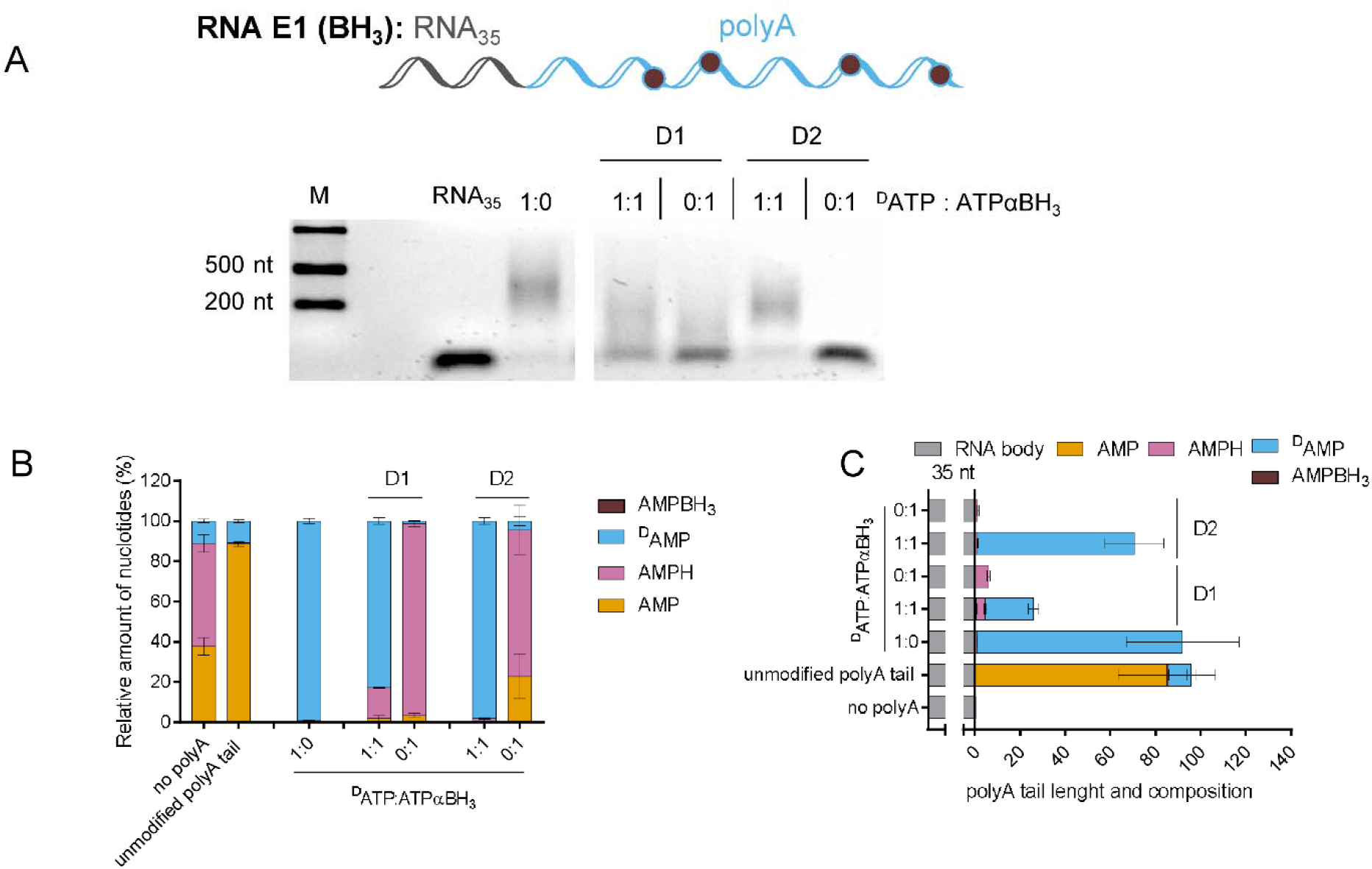
Boranophosphate RNAs release adenosine *H*-phosphonate (AMPH) as a degradation product. Analysis of short RNAs (RNA E1) (with U instead of A in the sequence) with polyA tails added by polyA polymerase (PAP) using different DATP/ATP analog ratios. ^D^ATP includes 10% ATP, which is taken into account during data analysis. RNA E1 is digested using snake venom phosphodiesterase (SVPDE) and Ccr4-Not transcription complex subunit 7 (CNOT7) for 1 h at 37 □C (see Materials and Methods for further details). (A) Data shown are from a single agarose gel. (B) Amount of nucleotides (the mean ± standard error of the mean (SEM)) after RNA degradation, as determined by liquid chromatography– tandem mass spectrometry (LC-MS/MS) normalized to guanosine monophosphate (GMP); (C) PolyA tail length (the mean ± SEM), calculated based on the concentration of nucleotides released from the RNA and RNA body sequence (Materials and Methods).

### 2.7. Boranophosphate modifications had a moderate influence on polyA tail susceptibility to deadenylation and diminished protein expression levels in cells

To study translational properties of mRNAs with boranophosphate-modified polyA tails, IVT RNA H was obtained by transcription *in vitro,* purified by RP HPLC (Figure S7C), and polyadenylated with PAP in the presence of ATP, ATPαBH_3_ D1 or ATP:ATPαBH_3_ D1 mixtures (at 49:1, 24:1, 14:1, 9:1, 5:1, 3:1, and 1:1 molar ratios) (Figure 8A). RP HPLC-purified RNA I, carrying a template-encoded polyA tail, was used as a reference in this experiment. The initial experiments revealed that the increased concentration of ATPαBH_3_ D1 in the reaction mix resulted in shorter polyA tails (Figure S8C), which was consistent with the results obtained for RNA E polyadenylated with ATPαBH_3_ D1 (Figure 7). To obtain RNA H1 variants with polyA tails of similar lengths, the polyadenylation protocol (previously set up for RNA G) was modified by adjusting the incubation time with PAP. Using this approach, we obtained RNA H1 variants with ~200– 250 nt polyA tails (based on an agarose gel). The obtained RNA H1 variants were highly homogenous (Figures S13-S15), regardless of the ATP:ATPαBH_3_ D1 ratio used for polyadenylation; therefore, we also prepared an RNA H1 variant with a polyA tail synthesized in the presence of 100% ATPαBH_3_ D1 (Figures S13, S14).

**Figure 8.**
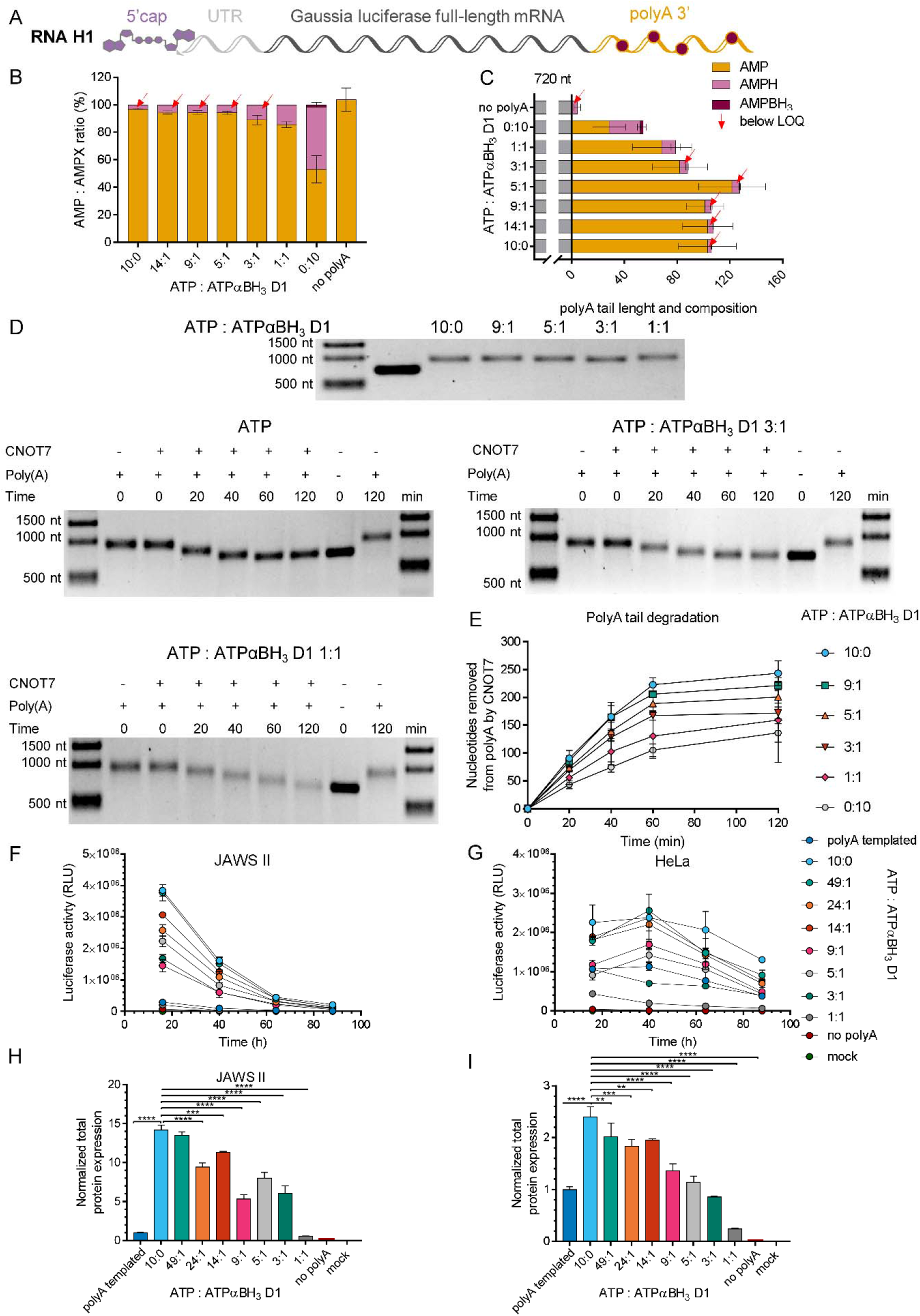
Analysis of mRNAs carrying boranophosphate moieties within the polyA tail. (A) Schematic representation of the *Gaussia* luciferase-coding transcript, RP HPLC-purified and polyadenylated with polyA polymerase (PAP) in the presence of ATP, ATPαBH_3_ D1 alone, or various mixtures of ATP:ATPαBH_3_ D1 (14:1,9:1, 5:1, 3:1, or 1:1). (B) RNA H1 tails composition (the mean ± standard error of the mean (SEM)) and (C) average polyA tail lengths (the mean ± SEM) obtained from LC-MS/MS. (D) Analysis of RNA H1 susceptibility to deadenylation. RNA H1 variants are incubated with CNOT7 deadenylase for 0–120 min, and the RNA length is analyzed on a 1% agarose gel using Image Lab 6.0.1 Software (Bio-Rad). The length of the polyA tail at each time point is estimated as the difference between digested RNA H1 and RNA H (transcript before polyadenylation). (E) Susceptibility to deadenylation of RNA H1 polyadenylated with various ATP:ATPαBH_3_ D1 ratios, estimated as the number of nucleosides removed by CNOT7 deadenylase at particular time points. (F-I) Translational properties of RNA H1 with boranophosphate-modified polyA tails. *Gaussia* luciferase activity in the supernatant of (F) JAWS II and (G) HeLa cells, as measured 16, 40, 64, and 88 h after transfection with RNA H1. The cell medium was exchanged after each measurement. Data points represent mean values ± standard deviation (SD) of one biological replicate consisting of three independent transfections. (H) Total protein expression (cumulative luminescence) over 4 days in JAWS II and (I) HeLa cells calculated from the same experiment. Bars represent the mean value ± SD normalized to RNA I (with a template-encoded A_128_ polyA tail). Statistical significance: * *p* < 0.05, ** *p* < 0.01, *** *p* < 0.001, **** *p* < 0.0001 (one-way analysis of variance (ANOVA) with Dunnett’s multiple comparison test). Only statistically significant differences are marked on the graph.

To determine the composition and average polyA tails lengths, different variants of RNA H1 were enzymatically degraded with SVPDE and CNOT7 ribonucleases, and the mixtures of the obtained NMPs were analyzed using LC-MS/MS (Figure 8B,C). The determined frequencies for ATPαBH_3_ D1 were lower than that for ATPαS D1, especially for RNA H1 variants polyadenylated by PAP at high concentrations of ATPαBH_3_ D1 (Figure 6B, Figure 8B, Table S4).

To evaluate the influence of boranophosphate modification on the polyA tail susceptibility to deadenylation, the transcripts were subjected to degradation by CNOT7 and the reaction progress was monitored using agarose gel electrophoresis. Based on the determined polyA tail lengths as a function of time, deadenylation rates were determined for each RNA (Figure 8E). The degradation rate for unmodified RNA H1 was similar to the previously analyzed unmodified RNA G1 (V_0_ 4.1 nt/min and V_0_ 4.3 nt/min, respectively) (Table S6, S7), suggesting that the RNA body sequence had no influence on susceptibility to CNOT7. RNA H1, with a boranophosphate-modified polyA tail, was more susceptible to deadenylation by CNOT7 than its counterparts containing phosphorothioate moieties at similar frequencies (Figure 6D,E, Figure 8D,E, Table S4). Interestingly, RNA H1 polyadenylated with 100% ATPαBH_3_ D1 was degraded faster than RNA G1 polyadenylated by PAP at a 9:1 molar ratio of ATP:ATPαS D1 (according to LC-MS/MS, the phosphorothioate moiety content was 6%). These results showed that phosphorothioate modification had greater potential to affect the deadenylation process than a boranophosphate moiety.

To analyze the translational properties of RNA H1 polyadenylated with selected ATP:ATPαBH_3_ D1 molar ratios, JAWS II and HeLa cells were transfected with RNA H1 variants and RNA I, and the activity of *Gaussia* luciferase secreted into the medium was measured over time (Figure 8F,G). It was found that total protein expression levels decreased with increasing boranophosphate content (Figure 8H,I). Even for the lowest tested ATP:ATPαBH_3_ D1 ratio (49:1), the translational efficiency was slightly compromised. This result suggested that the boranophosphate modification was less compatible with mRNA expression *in vivo* than phosphorothioate.

## 3. Discussion

In this work we contributed to the development of chemically-modified mRNAs with superior biological properties. So far, properties of therapeutic mRNAs were modulated by the use of chemical 5’ cap analogs, introducing unnatural nucleosides into the mRNA body, switching between various 5’ and 3’ UTR elements, or by optimizing codons. It has been previously demonstrated that replacing one of the four nucleotides used for *in vitro* transcription by its phosphorothioate analog produces mRNAs that are efficiently expressed in a prokaryotic cell-free translation system (72). However, prokaryotic mRNAs do not contain the 5’ and 3’ end stabilizing elements (5’ cap and polyA tail, respectively), characteristic for eukaryotic mRNAs, and are processed by quite different translational machineries(77), so the biological consequences of phosphate-modifications may be different for these two systems. We have previously shown that an O-to-S or O-to-BH_3_ substitution within the β-phosphate of the mRNA cap confers beneficial translational properties to mRNA by stabilizing the interaction with eIF4E and blocking 5’→3’ degradation (10,78). To the best of our knowledge, the presence of O-to-X substitutions within the eukaryotic mRNA body has not been studied. Here, we investigated whether the presence of O-to-S or O-to-BH_3_ substitutions within the 5’,3’-phosphodiester bonds of mRNA influenced translational properties in mammalian expression systems. We envisaged that such modifications may have beneficial effects on mRNA stability, primarily by blocking the degradation from the 3’ end (the polyA tail). To test this hypothesis, we first developed a method for quantitative assessment of incorporation of ATP analogs modified at the α-phosphate into RNA by polymerases, which relied on exhaustive degradation of the transcripts by 3’ nucleases followed by LC-MS/MS analysis of the resulting nucleoside monophosphates and their phosphate modified analogs. The phosphorothioate-containing RNAs released AMPS as the main degradation product, whereas the boranophosphate RNAs, released AMPH instead of AMPBH_3_, which was unexpected, but did not hamper method development. The key factor during method optimization was the adjustment of degradation conditions to ensure complete RNA degradation without desulfurization of AMPS to AMP. The method was successfully used to analyze both short and long RNAs, with the phosphorothioate quantification limit found to be 0.03 μM. The incorporation assessment of ATPαS diastereomers (D1 = *S*_P_ and D2 = *R_P_)* by SP6 and T7 polymerases, revealed that only the *S*_P_ isomer of ATPαS served as the unnatural substrate for these polymerases, albeit not as efficiently as ATP, which was most evident from the decreased transcription yield when ATPαS D1 was used as the only adenine-containing substrate (Figure 3, 4, S4). The *R*_P_ isomer of ATPαS was neither substrate nor inhibitor of the polymerases. These findings are in good agreement with literature reports on stereo-preferences of RNA polymerases (65,79–83). Degradation of RNAs obtained with a deuterated ATP analog (^D^ATP) and either ATPαS D1 or ATPαBH_3_ D1 at a 1:1 ratio under optimized conditions revealed less than 10% of non-deuterated AMP (~7% for phosphorothioate and 2% for boranophosphate), indicating that more than 90% of the modified analyte remained intact in the presence of 5’ nucleases (Figure 7, S5). Following these initial experiments, we set out to evaluate biological properties of mRNAs IVT in the presence of ATP analogs. We found that in the presence of ATPαS D1: ATP mixtures, IVT mRNAs containing template-encoded polyA tails were synthesized significantly less efficiently by both T7 and SP6 polymerase than in the presence of ATP, which limited the amount of mRNA available to further biochemical analyses. Moreover, the yields of protein expression (firefly luciferase) per nanogram of RNA measured in RRLs were also significantly lower than those for unmodified mRNAs. This is in contrast to prokaryotic systems, where the detrimental effect of mRNA body modification within phosphorothioate groups on translation of prokaryotic mRNA has been reported only if more than one NTP was replaced with NTPαS (72). It should be noted, however, that the firefly luciferase-based assay used in our work detected only functional proteins, so shortened, misfolded or mutated proteins may be formed, but were not detected. We hypothesized that phosphorothioate modifications within the UTR and ORF may impair ribose movement necessary for efficient translation. Therefore, we focused on modifying mRNA exclusively within the polyA tails. Modified polyA tails were introduced into IVT mRNA by the use of PAP from *E.coli* as a specific tailing enzyme. The same enzyme has been recently used to fluorescently label polyadenylated mRNA (33), but to the best of our knowledge, it has never been studied with phosphate-modified ATP analogs. The polyA tail lengths estimated by LC-MS/MS were slightly shorter than those obtained with ATP under the same conditions. However, the data on the length should be interpreted carefully due to differences in RNA homogeneity observed for different RNA types and compositions. The fact that polyA tails obtained in the presence of PAP are highly heterogeneous has been previously reported (84,85). The incorporation of ATPαX D1 analogs was even less efficient in the absence of ATP.

The polyA tail impacts mRNA stability and translation efficiency via interaction with PABPs (86,87). Recent studies highlight a correlation between polyA tail length and translation of mRNA (22). Therefore, to analyze P-mod mRNA susceptibility to deadenylation *in vitro* and expression in living cells, we prepared mRNAs of similar polyA tails lengths (200–250 nucleotides long) (88). This was confirmed using both electrophoretic analysis and LC-MS/MS (Figure 6, 8). The susceptibility of mRNAs with phosphate-modified polyA tails to deadenylation by recombinant human CNOT7, which is one of the two catalytic subunits of Ccr4-Not complex(89), was analyzed by gel electrophoresis (Figure 6, 8). We observed a significantly decreased susceptibility of mRNAs polyadenylated by PAP to deadenylation in the presence of ATP:ATPαS D1 mixtures. The degradation rates correlated qualitatively with the modification frequency (the higher the phosphorothioate content, the more stable the polyA). In contrast, boranophosphate modification only moderately affected susceptibility to CNOT7; the boranophosphate mRNA was ~2.7-fold more susceptible to CNOT7 than a phosphorothioate mRNA of analogous composition. As the structural information elucidating the atomic details of the polyA recognition by CNOT7 deadenylase is not available, it is difficult to speculate on the different stabilities of phosphorothioate and boranophosphate RNAs. Nonetheless, the differences in size, charge distribution, and solvation between a sulfur atom and a borano group affect hydrogen-bonding, metal ion coordination potential (90,91), and polyA topology, which in turn leads to various rearrangements of the CNOT7 active site upon substrate binding and differentially affects catalytic efficiency. The differentiation between these two modifications by CNOT7 may guide a better understanding of substrate specificity of deadenylases, and hence phosphate-modified polyA tails may serve as useful research tools.

To investigate whether polyA stability *in vitro* impacts protein biosynthesis in living cells, we measured the expression of *Gaussia* luciferase-encoding mRNAs containing variously modified polyA tails in two cell lines: HeLa (cancer cells) and JAWS II (dendritic cells, relevant in the context of mRNA-based vaccines). In both cell lines, phosphorothioate-modified polyA tails did not affect protein expression, regardless of the phosphorothioate moiety content in the polyA tail. This indicated that the phosphorothioate modification was compatible with translation in living cells, albeit it did not provide any evident biological effect, at least for the studied polyA tail compositions. In contrast, the presence of a boranophosphate moiety in polyA tails clearly correlated with a decline in protein expression levels. Overall, the data from both *in vitro* and cell culture experiments indicated that only the phosphorothioate modification stabilized the polyA tail and efficiently supported translation. It remains to be determined how this modification affects interaction with PABPs. As demonstrated by Deo et al. (92) and Safaee et al. (93), PABP-oligoA_11_ complex formation is mostly driven by extensive interactions of PABPs with adenines, engaging hydrogen bonds, van der Waals contacts and base stacking. The sugar-phosphate backbone is less extensively probed by PABP amino acid side chains; therefore, potentially providing steric freedom for substitutions of the non-bridging oxygen atoms of the α-phosphate (92,93). Nevertheless, phosphorothioate and boranophosphate modifications can affect three-dimensional organization of the polyA tail; therefore, influencing the dynamics of its recognition by PABPs (94,95). Finally, it should be emphasized that the polyA tails obtained by the reported enzymatic procedure were heterogeneous not only in length, but also in composition. Random incorporation of ATPαS resulted in a mixture of thousands of polyA variants in one synthesis batch, which were inseparable and thus analyzed together as one sample. However, it is likely that these variants might quite significantly differ in biological properties. The fact that the average translation for these variants was comparable to unmodified mRNA, did not exclude the existence of modification “sweet-spots” characterized by superior biological properties. Although currently available synthetic methods do not allow for fully-controllable incorporation of phosphate modification into polyA, our results indicate that this field is worth further exploration in the context of therapeutic mRNA.

Overall, our research is the first step towards understanding the impact of sugar-phosphate mRNA backbone modifications on the biological properties of these molecules. The mRNA synthesis and analysis protocols reported here may find application in basic studies dissecting mRNA translation and degradation, as well as studying properties of mRNA-based therapeutics. Intriguingly, phosphorothioate modification has been recently identified in rRNA in *E. coli, L. lactis,* and HeLa cells, which might spike even higher interest in the properties of phosphate-modified RNAs in the future (96).

## 4. Experimental Section

### 4.1. Chromatographic and mass spectrometric conditions

#### 4.1.1. Instrumentation

All LC-MS/MS analyses were performed on a QTRAP 3200 (AB Sciex) system consisting of an electrospray ion source and triple quadrupole/linear ion trap analyzer. The instrument was coupled with an Agilent Technologies high performance liquid chromatography (HPLC) system equipped with 1260 Bin Pump and degasser, 1260 ALS autosampler and 260 TCC column oven. Analyst 1.6 software was used for system control, data acquisition, and data processing.

#### 4.1.2. General information

MS-grade reagents (methanol, acetonitrile, ammonium acetate and other solvents) and starting materials (such as unmodified nucleosides and ATP) and chemical reagents, including N,N-dimethylhexylamine (DMH) and 97% H_2_^18^O were purchased from commercial sources. Nucleotide analogs were synthesized by published methods (53–55). SP6 and T7 polymerases were purchased from ThermoFisher Scientific. PAP was purchased from Lucigen. Snake venom phosphodiesterase (SVPDE) was purchased from Sigma Aldrich. Ccr4-Not transcription complex subunit 7 (CNOT7) was purified as described in the Materials and Methods.

If necessary, the compounds were purified by means of analytical HPLC or semi-preparative reversed-phase (RP) HPLC. Analytical HPLC (Series 1200; Agilent Technology) was performed using a Supelcosil LC-18-T HPLC column (4.6 × 250 mm, 5 μm, flow rate 1.3 mL/min) with a linear gradient of 0–50% methanol in 0.05 M ammonium acetate buffer (pH 5.9) for 15 min and UV detection at 254 nm. Semipreparative HPLC was performed on the same apparatus equipped with a Discovery RP Amide C-16 HPLC column (25 cm × 21.2 mm, 5 μm, flow rate 5.0 mL/min) with a linear gradient of 0–100% acetonitrile in 0.05 M ammonium acetate buffer (pH 5.9) for 120 min with UV detection at 260 nm.

#### 4.1.3. Liquid chromatography

Chromatographic separation of nucleoside 5’-monophosphates (NMPs) was achieved on an Eclipse XDB-C18 analytical column (5.0 μm, 4.6 mm × 150 mm, Agilent) equipped with an Eclipse XDB-C18 analytical guard column (5.0 μm, 4.6 × 12.5 mm). The gradient mobile phase contained the DMH ion pair reagent (56) and comprised two eluents: eluent A, which was 20 mM aqueous DMH adjusted to pH 4.8 with formic acid, and eluent B, which was a 1:1 (v/v) mixture of acetonitrile and aqueous 20 mM DMH adjusted to pH 4.8 with formic acid. The elution was performed at room temperature (RT) and at a flow rate of 700 μL min^-^. The gradient was optimized to obtain sufficient chromatographic separation of all analytes of interest (AMP, AMPS, and GMP). The optimal gradient was 0–100% eluent B in 15 min.

#### 4.1.4. Mass spectrometry

The analytes in the HPLC eluate were monitored by MS in multiple reaction monitoring (MRM) mode. MS conditions for individual analytes and internal standards were optimized directly from the syringe pump. The optimization was performed using 400 μM solutions of individual NMPs and their respective isotope-labeled internal standards. Optimizations and analyses have been performed in negative ionization mode to maximize the number of fragmentation reactions (57). The general ion source conditions used in all experiments were as follows: turbo ion-spray voltage:-4500 V, temperature: 300 °C, curtain gas: 30 psi, ion source gas 1: 30 psi, and ion source gas 2: 25 psi. The MRM transitions and MS conditions for individual NMPs and isotope-labeled internal standards are depicted in Table S1.

### 4.2. Synthesis of reference compounds and internal standards

#### 4.2.1. Nucleoside 5’-O-monophosphates: AMP, [^18^O,^18^O]AMP, GMP, [^18^O,^18^O]GMP

Nucleoside 5’-phosphorylation was essentially performed as described earlier (58), (Figure S1A). Briefly, an appropriate nucleoside (0.01 mmol) was dissolved/suspended in trimethyl phosphate (0.5 mL) at 0 °C followed by the addition of phosphoryl chloride (POCl_3_, 0.03 mmol). The reaction was monitored by RP HPLC (as described above in General information). The reaction was quenched with either 97% [^18^O]-H_2_O or H_2_O, when the conversion of the substrate reached 60–90%. pH was neutralized using NaHCO_3_.The desired nucleoside 5’-monophosphate was purified by RP HPLC. The purity and isotopic composition of the isolated products were analyzed by electrospray ionization (ESI,-)/MS (Table S2). The analyses confirmed that the products obtained via hydrolysis with heavy water contained at least 87% of the expected [^18^O,^18^O] double-labeled nucleoside 5’-monophosphate along with smaller amounts (12%) of [^18^O] single-labeled product, and less than 1% of unlabeled product.

#### 4.2.2. Adenosine 5’-O-monothiophosphate: AMPS, [^18^O,^18^O]AMPS

Nucleoside 5’-thiophosphorylation was essentially performed as described earlier (59), (Figure S1A). Briefly, adenosine (0.01 mmol) was suspended in trimethyl phosphate (0.5 mL) and placed in an ice-bath for 5 min, followed by addition of 2,6-lutidine (0.04 mmol) and thiophosphoryl chloride (PSCl_3_, 0.03 mmol). The reaction progress was monitored by RP HPLC. The reaction was quenched after 6 h with either 97% [^18^O]-H_2_O or H_2_O, when conversion of the substrate reached 60–90%. pH was neutralized using NaHCO_3_.The products were purified by RP HPLC (as described above in General information). The purity and isotopic composition of the isolated products was analyzed by ESI(-)/MS (Table S2). The analyses confirmed that the products obtained via hydrolysis with heavy water contained at least 87% of the expected [^18^O,^18^O] double-labeled nucleoside 5’-monophosphate along with smaller amounts (11%) of [^18^O] single-labeled product, and less than 2% of unlabeled product.

#### 4.2.3. Adenosine-5’-O-(*W-*phosphonate): AMPH, [^18^O,^18^O] AMPH

AMPH and [^18^O,^18^O] AMPH were synthesized as previously described (10) (Figure S1B). 2’,3’-O-isopropylidene-adenosine (0.65 mmol) was suspended in trimethyl phosphate (3 mL) and placed in an icebath followed by the addition of phosphorus trichloride (PCl_3_, 1.95 mmol). Reaction progress was monitored by RP HPLC and quenched with 97% [^18^O]-H_2_O or H_2_O after 0.5 h. Then, sodium bicarbonate (NaHCO_3_) was added to neutralize the pH, and the solution was kept for 6 h at RT to remove the isopropylidene protecting group. The product was purified by HPLC (as described above in General information). The purity and isotopic composition of the isolated products was analyzed by ESI(-)/MS (Table S2). The analyses confirmed that the products obtained via hydrolysis with heavy water contained at least 84% of the expected [^18^O,^18^O] double-labeled nucleoside 5’-monophosphate along with smaller amounts (10%) of [^18^O] single-labeled product, and less than 6% of unlabeled product.

#### 4.2.4. Adenosine 5’-O-monoboranophosphate: AMPBH_3_, [^18^O,^18^O] AMPBH_3_

AMPBH_3_ and [^18^O,^18^O] AMPBH_3_ were obtained as previously described (10). Adenosine-5’-O-(*H*-phosphonate) or [^18^O,^18^O] adenosine-5’-O-(*H-*phosphonate) (0.06 mmol) was placed in a rubber septum capped round-bottom flask under argon atmosphere and suspended in acetonitrile (1 mL). Then, N,O-bis(trimethylsilyl)acetamide (BSA, 1.2 mmol) was added through a syringe and after 30 min of stirring, BH3·SMe_2_ (0.12 mmol) was added and the mixture was left for 30 min. An adequate product was released after addition of a trimethylamine (TEA)/methanol solution. The solution was evaporated to dryness and three times re-evaporated from methanol. Products were purified by HPLC (as described above in General information). The purity and isotopic composition of the isolated products was analyzed by ESI(-)/MS (Table S2). The analyses confirmed that the products obtained via hydrolysis with heavy water contained at least 99% of the expected [^18^O,^18^O] double-labeled nucleoside 5’-monophosphate.

#### 4.2.5. 8-Deuteroadenosine 5’-O-triphosphate: ^D^ATP

Adenosine 5’-*O*-triphosphate sodium salt (3 mg) was diluted in 1 mL of D_2_O and TEA was added to pH 10. The mixture was incubated at 60 °C. The progress of gradual H-to-D exchange was monitored by HPLC-MS. After 2 d, the sample was freeze-dried, redissolved in D_2_O in the presence of TEA, and the incubation was continued at 60 °C. The procedure was repeated two more times (after 2 and 4 days). After 6 d total incubation time, the exchange efficiency reached 94%. The product was purified by HPLC and freeze-dried three times.

### 4.3. Method validation

#### 4.3.1. Calibration curves

The calibration curves were prepared by spiking known concentrations (0.01–50 μM) of NMP and a constant concentration of isotopically labeled internal standard (2 μL, 10 μM; the final concentration in 20 μL samples was 1 μM) into reaction solution (SVPDE buffer (1 mM magnesium chloride (MgCl_2_), 5 mM Tris, pH 8,8) and deactivated SVPDE) (Figure S2). Calibration curves were built by fitting the analyte-to-internal standard concentration ratio versus the analyte-to-internal standard peak area ratio using linear regression analysis *y=ax+b,* where the *a* parameter is the slope and *b* is the y-intercept. The limits of quantification are depicted in Table S3.

#### 4.3.2. Degradation conditions

A typical RNA degradation reaction (20 μL) was performed at 37 °C for 1 h and contained: either 1) 20 ng RNA_35_ (RNA A, RNA B and RNA C), 3000 ng/mL SVPDE (Sigma Aldrich) in SVPDE buffer (1 mM MgCl_2_, 5 mM Tris, pH 8,8); or 2) 20 ng RNA_35_ with polyA (RNA A1 and RNA E1), or 40 ng (RNA F2 and RNA G1) (or more ~120 ng, in the case of the boranophosphate modification, RNA H) mRNA, 9000 ng/mL SVPDE (Sigma Aldrich), 1 mg/mL CNOT7 enzyme, buffer (5 mM MgCl_2_, 10 mM Tris, pH 8.0, 50 mM potassium chloride (KCl), 10 mM dithiothreitol (DTT)). The reactions were quenched by snap-freezing in liquid nitrogen, followed by heating at 95 °C for 5 min, and snap-freezing in liquid nitrogen again. Samples after degradation were analyzed using LC-MS/MS. To each sample containing 20 ng/40 ng/120 ng of degraded RNA, 1 μM isotopically labeled internal standard was added. The enzyme amount was initially optimized on model RNAs to ensure degradation to single mononucleotides and lack of their further degradation (which was observed by MS at high enzyme concentrations, Figure 2F,G Figure S3). During optimization, the post-degradation reaction mixtures were analyzed by 15% polyacrylamide gel electrophoresis (PAGE). If any undigested RNA fragments were noticeable, the enzyme amount was increased.

#### 4.3.3. Solid-phase synthesis of oligoribonucleotide 5’-phosphate pG(CU)_5_A_11_ used as a control sample (RNA C, Table 1)

Solid-phase synthesis of oligoribonucleotide was performed on a 1 μmol scale using AKTA Oligopilot plus 10 synthesizer (GE Healthcare) and Custom PrimerSupport™ Ribo A 40 (GE Healthcare) solid support. In the coupling step, 20 equivalents of N6-Benzoyl-2’-O-tert-butyldimethylsilyl-5’-O-DMT-adenosine 3’-CE (2’-*O*-TBDMS) phosphoramidate (rA^Ac^, rC^Ac^, rG^Ac^, U or biscyanoethyl phosphoramidite, all from ChemGenes) and 0.30 M 5-(benzylthio)-1-*H*-tetrazole in acetonitrile were recirculated through the column for 15 minutes. A solution of 3% (v/v) dichloroacetic acid in toluene was used as a detritilation reagent, 0.05 M iodine in pyridine/water (9:1) for oxidation, 20% (v/v) *N*-methylimidazole in acetonitrile as Cap A and a mixture of 40% (v/v) acetic anhydride and 40% (v/v) pyridine in acetonitrile as Cap B. After the last cycle of synthesis, RNA, still on the solid support, was treated with 20% (v/v) diethylamine in acetonitrile to remove the 2-cyanoethyl protecting groups. The product was cleaved from the solid support and deprotected with AMA (methylamine/ammonium hydroxide 1:1_v/v_; 55 °C, 1 h), evaporated to dryness and redissolved in dimethyl sulfoxide (100 μL). The TBDMS groups were removed using triethylammonium trihydrofluoride (TEA·3HF; 125 μL, 65 °C, 3 h) and the oligonucleotide was desalted by precipitation with sodium acetate and 1-butanol. Finally, the product was purified by RP HPLC (linear gradient of acetonitrile in ammonium acetate (CH_3_COONH_4_, pH 5.9) to give, after repeated freeze-drying from water, an ammonium salt of pG(CU)_5_A_11_. For LC-MS/MS analysis, 20 ng oligoRNA was digested using SVPDE, according to the above conditions.

### 4.4. Calculations – concentrations and polyA tail length

Concentrations of analytes (*C_A_*) were calculated based on calibration curve parameters *(a,b),* internal standard concentration (*C_IS_*), and signal intensities (*I_A_* – analyte intensity, *I_IS_* – internal standard intensity); *b* was fixed to 0.0 (Equation 1).

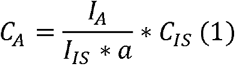

Concentrations of AMP in the polyA tail were calculated according to Equation 2.

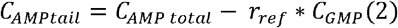

*r_ref_* is a value obtained from the appropriate RNA fragment without the polyA and is the ratio of the concentration of AMP and GMP 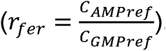. Numbers of adenine nucleotides in the polyA tail (*n_AMptail_*) were calculated based on Equation 3 and numbers of G in a sequence (*n_CMPref_*):

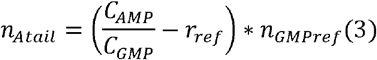

Numbers of adenine nucleotide analogs *(n_AMPXtail_)* were calculated according to Equation 4:

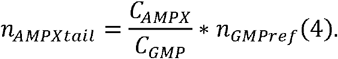

### 4.5. RNA synthesis and purification

The sequences of RNAs used in different types of experiments are summarized in Table 1. The preparations for each RNA are described below.

**Table 1.**
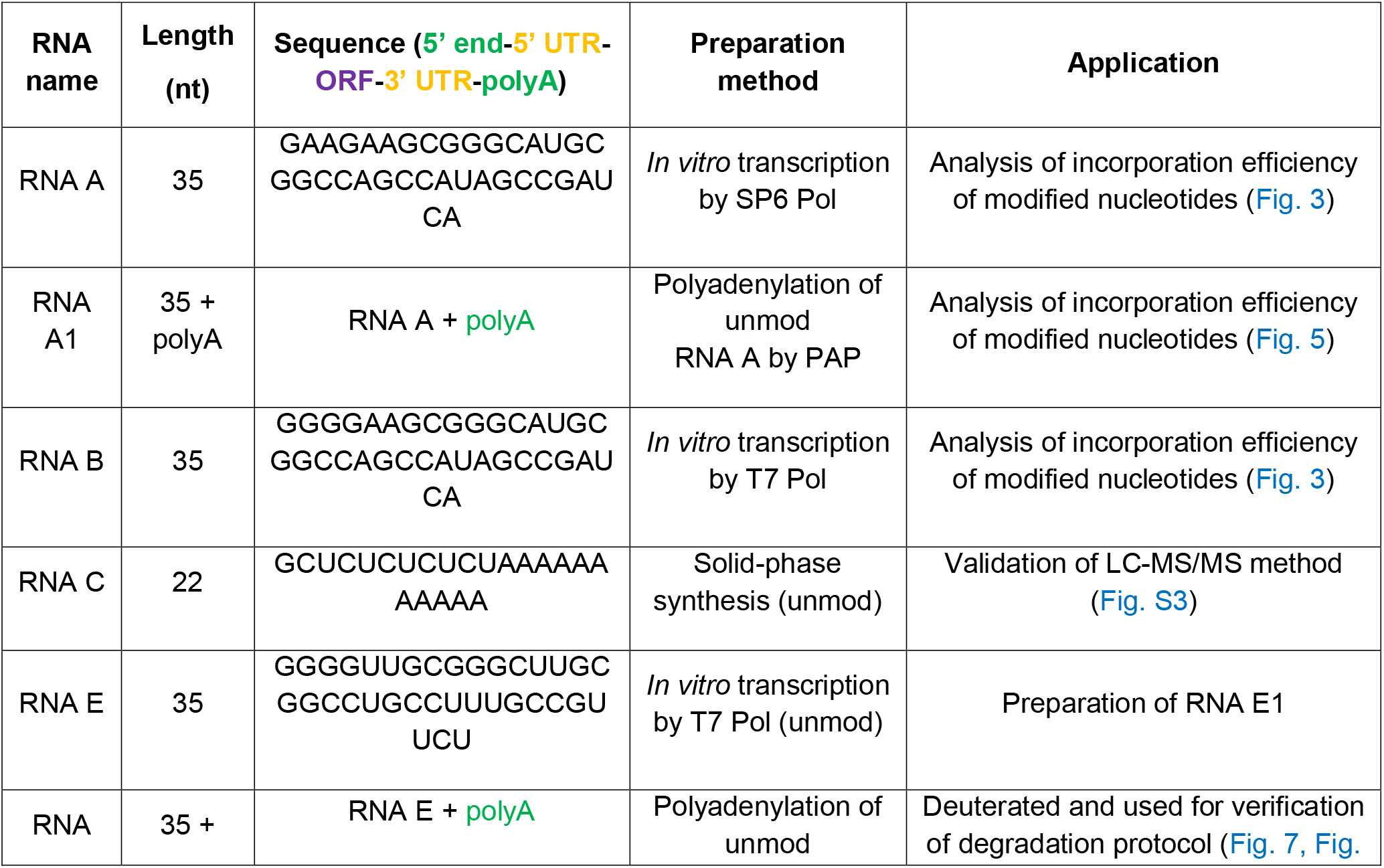
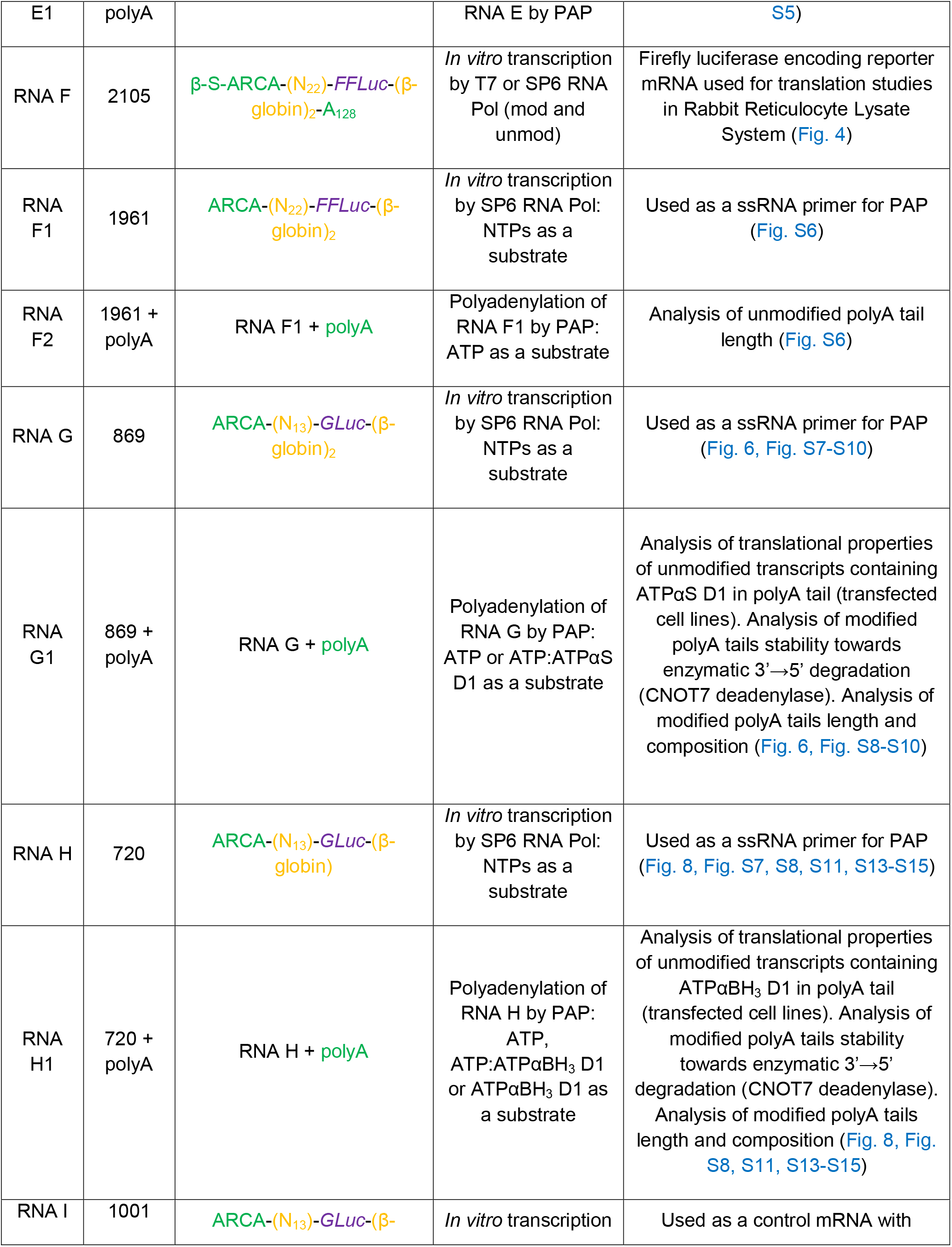
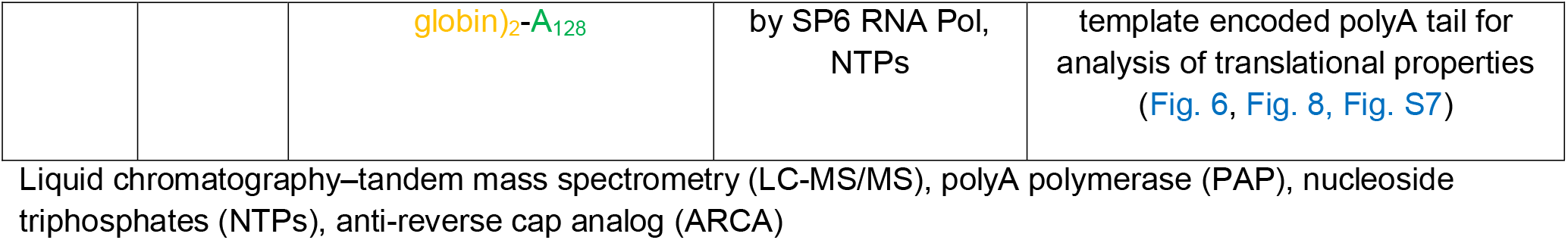
RNA sequences studied in this work

#### 4.5.1. Short RNA (RNA A and B)

Short RNAs were generated on a template of annealed oligonucleotides: 1) (CAGTAATACGACTCACTATAGGGGAAGCGGGCATGCGGCCAGCCATAGCCGATCA and TGATCGGCTATGGCTGGCCGCATGCCCGCTTCCCCTATAGTGAGTCGTATTACTG) (60), which contains the Φ6.5 T7 promoter sequence (TAATACGACTCACTATA) and encodes a 35-nt long sequence (GGGGAAGCGGGCATGCGGCCAGCCATAGCCGATCA); 2) (ATACGATTTAGGTGACACTATAGAAGAAGCGGGCATGCGGCCAGCCATAGCCGATCA and TGATCGGCTATGGCTGGCCGCATGCCCGCTTCTTCTATAGTGTCACCTAAATCGTAT) (60), which contains the SP6 promoter sequence (CGA TTTAGG TGACAC TATA) and encodes a 35-nt long sequence (GA AGAAGC GGGCAT GCGGCC AGCCAT AGCCGA TCA). ATP analogs used for RNA preparation were synthesized, as previously described (10,54). A typical *in vitro* transcription reaction (20 μL) was incubated at 37 °C for 2 h and contained: RNA polymerase buffer (40 mM Tris-HCl pH 7.9, 6 mM MgCl_2_, 1 mM DTT, and 2 mM spermidine), 10 U/μL T7 or 1 U/μL SP6 RNA polymerase (ThermoFisher Scientific), 1 U/μL RiboLock RNase Inhibitor (ThermoFisher Scientific), 0.5 mM CTP/UTP/GTP, 0.5 mM different ATP:ATPαX ratios, and 0.4 μM annealed oligonucleotides as a template. Following a 2 h incubation, 1 U/μL DNase I (ThermoFisher Scientific) was added and the incubation was continued for 30 min at 37 °C. The crude RNAs were purified using RNA Clean & Concentrator-25 (Zymo Research). The quality of transcripts was assessed on 15% acrylamide/7 M urea gels, whereas the concentration was determined spectrophotometrically. To remove *in vitro* transcription side-products of unintended size, RNA samples were gel-purified using PAA elution buffer (0.3 M sodium acetate, 1 mM ethylenediaminetetraacetic acid (EDTA), and 0.05% Triton X-100), precipitated with isopropanol, and dissolved in water.

#### 4.5.2. Short RNA without an A in the sequence (RNA E)

Short RNAs were generated on a template of annealed oligonucleotides (the same sequence as RNA B but with U instead of A) (CAGTAATACGACTCACTATAGGGGTTGCGGGCTTGCGGCCTGCCTTTGCCGTTCT and AGAACGGCAAAGGCAGGCCGCAAGCCCGCAACCCCTATAGTGAGTCGTATTACTG), which contained the T7 promoter sequence (TAATACGACTCACTATA) and encoded a 35-nt long sequence (GGGGTTGCGGGCTTGCGGCCTGCCTTTGCCGTTCT). The *in vitro* transcription reaction (200 μL) was incubated at 37 °C for 3 h and contained: RNA polymerase buffer (40 mM Tris-HCl pH 7.9, 20 mM MgCl_2_, 1 mM DTT, and 2 mM spermidine), 10 U/μL T7 RNA polymerase (ThermoFisher Scientific), 1 U/μL RiboLock RNase Inhibitor (ThermoFisher Scientific), 5 mM CTP/UTP/GTP, and 1 μM annealed oligonucleotides as a template. Following 3 h incubation, 1 U/μL DNase I (ThermoFisher Scientific) was added and the incubation was continued for 30 min at 37 °C. IVT products were extracted using phenolchloroform and purified using HPLC on a Clarity 3 μm Oligo-RP C18 column (Phenomenex). To elute the desired products, two eluents were used: Eluent A: 50 mM TEA and Eluent B: 75% acetonitrile in the following gradient: 0–5 min 95% A, 5–20 min 87.5% A, 20–21 min 50% A using a 1 mL/min flow rate. Obtained products were used to obtain short RNAs with a polyA tail (RNA E1), as described below.

#### 4.5.3. Short RNA with the polyA (RNA A1 and E1)

Short RNAs with a polyA tail were generated in a polyadenylation reaction. A typical reaction mixture (20 μL) was incubated at 37 °C for 1 h and contained: 1 μg 35 nt RNA fragment, 1 mM different ATP([D]ATP):ATPαX ratios, 1 U/μL RiboLock RNase Inhibitor (ThermoFisher Scientific), PAP dedicated buffer and PAP (Lucigen) from 0.05 to 0.2 U/μL reaction. The RNAs were purified using RNA Clean & Concentrator-25 (Zymo Research). The quality of transcripts was assessed on 2% agarose gels, whereas the concentration was determined spectrophotometrically.

#### 4.5.4. *Firefly* luciferase (RNA F and F1)

For studies of transcription and translation efficiency of mRNA uniformly modified with phosphorothioate, *Firefly* luciferase transcripts (RNA F) were prepared based on the pJET_T7_FFLuc_128A and pJET_SP6_FFLuc_128A plasmids (61). The DNA template was purified with GeneJET Plasmid Midiprep Kit (ThermoFisher Scientific) and linearized with the AarI restriction enzyme (ThermoFisher Scientific). The restriction site for AarI was located downstream of the polyA tract, therefore, IVT RNA F contained a template-encoded polyA tail. The completeness of plasmid linearization was evaluated by 1% 1× Tris-boric acid-EDTA (TBE) agarose gel electrophoresis, the sample was purified with QIAquick PCR Purification Kit (Qiagen) and used as a DNA template for *in vitro* transcription. The *in vitro* transcription reaction mix (30 μL) was maintained for 4 h at 40 °C and contained: transcription buffer (40 mM Tris-HCl pH 7.9, 10 mM MgCl_2_, 1 mM DTT, and 2 mM spermidine), 0.33 U/μL SP6 or T7 RNA polymerase, 0.66 U/μL RiboLock RNase inhibitor, 0.5 mM ATP or 0.5 mM mixture of ATP and ATPαS D1,0.5 mM CTP and UTP, 0.125 mM GTP, 1.25 mM β-S-ARCA (62) as a cap analog, and 120 ng of linearized plasmid. *In vitro* transcription was followed by removal of the DNA template with DNase I (30 min incubation at 37 °C with 0.06 U/μL of the enzyme), and purification with NucleoSpin RNA Clean-up XS (Macherey-Nagel). The quality of mRNA was analyzed using 1% 1× TBE agarose gel electrophoresis, and the concentration was determined spectrophotometrically (NanoDrop 2000c).

#### 4.5.5. *Gaussia* luciferase (RNA G, H, and I)

For analysis of polyA tail composition, length, stability and translational properties, three types of mRNA encoding *Gaussia* luciferase were prepared, differing in the number of *H. sapiens* β-globin 3’ UTR sequences and the presence or absence of a polyA tail: RNA I with a double 3’ UTR and template encoding a polyA tail, and RNA H and RNA G without polyA tails, and with single or double 3’ UTR sequences, respectively. The pJET_SP6_GLuc_128A plasmid (61), purified with the GeneJET Plasmid Midiprep Kit (ThermoFisher Scientific), was used as a DNA template for a polymerase chain reaction (PCR) performed using Phusion High-Fidelity DNA polymerase (ThermoFisher Scientific). Two amplification strategies were involved to obtain the template lacking an encoded polyA sequence (RNA G and H), and therefore suitable for further polyadenylation of IVT mRNA. The DNA template for *in vitro* transcription of RNA H was amplified using the SP6_Gluc_For_II (5’-GTCCCAATTAGTAGCATCACGCTGTG-3’) and pJET_Luc_FL_UTR_REV (5’-GCAATGAAAATAAATGTTTTTTATTAGGCAGAATCCAAATGC-3’) primers and was purified with the QIAquick PCR Purification Kit (Qiagen) directly after the PCR reaction. For preparation of the DNA template for *in vitro* transcription of RNA G and RNA I, the SP6_Gluc_For_II (5’-GTCCCAATTAGTAGCATCACGCTGTG-3’) and pJET_SQ_rev_II (5’-GCCAAGAAAACCCACGCCACCTAC-3’) primers were used. For the amplified DNA template, an additional step of restriction digestion was necessary to remove the sequence downstream of the second 3’ UTR (for *in vitro* transcription of RNA G) or downstream of the polyA coding region (for *in vitro* transcription of RNA I). To obtain highly concentrated DNA for more efficient restriction digestion, the amplification product was extracted from the PCR reaction mixture with phenol:chloroform (1:1, v/v) and precipitated with 1.25 M ammonium acetate and isopropanol at RT, followed by a washing step with 70% ethanol. The precipitated DNA pellet was resuspended in 20–50 μL RNase/DNase free water (Sigma). To obtain the DNA template for *in vitro* transcription of RNA G, restriction digestion using MssI (PmeI) (ThermoFisher Scientific) was performed, resulting in blunt-ended double-stranded DNA (dsDNA) containing a double 3’ UTR sequence and lacking a polyA tail. To prepare the DNA template for *in vitro* transcription of RNA I, the AarI (ThermoFisher Scientific) restriction enzyme was used. The restriction digestion efficiency was validated using 1% 1× TBE agarose gel electrophoresis and purified with a QIAquick PCR Purification Kit (Qiagen).

A typical *in vitro* transcription reaction mixture (30–40 μL) was incubated for 4 h at 40 °C and contained: transcription buffer (40 mM Tris-HCl pH 7.9, 10 mM MgCl_2_, 1 mM DTT, and 2 mM spermidine), 2.5 U/μL SP6 RNA polymerase (a second 2.5 U/μL was added after 2 h of incubation), 2 U/μL RiboLock RNase inhibitor, 2 mM ATP, CTP and UTP, 0.25 mM GTP, 2.5 mM ARCA, and 250 ng/μL DNA template. *In vitro* transcription was followed by removal of the DNA template with DNase I (30 min incubation at 37 °C with 0.1 U/μL of the enzyme). The crude transcript was initially purified with NucleoSpin RNA Clean-up XS (Macherey-Nagel), followed by a second purification step using RP HPLC. To isolate the homogenous mRNA fraction sufficient for further polyadenylation, obtained transcripts were purified on an Agilent Technologies Series 1200 HPLC using RNASep™ Prep (ADS Biotec) at 55 °C, as described previously (63). For mRNA purification, a linear gradient of buffer B (0.1 M triethylammonium acetate pH 7.0 and 25% acetonitrile) from 40–60% in buffer A (0.1 M triethylammonium acetate pH 7.0) over 25 min at 0.9 mL/min was applied. mRNA from collected fractions was recovered by precipitation with 0.3 M sodium acetate pH 5.2 and isopropanol at RT, followed by a washing step with 70% ethanol. The purity of mRNA was analyzed using 1% 1× TBE agarose gel electrophoresis and selected fractions were combined. The absorbance of the mRNA sample was measured spectrophotometrically (NanoDrop 2000c) and the concentration was determined using the extinction coefficient predicted for a particular mRNA sequence by DNA Calculator (software available on www.molbiotools.com website).

#### 4.5.6. mRNA polyadenylation by PAP (RNA G1 and H1)

The 3’ end of *Gaussia* mRNA transcripts was tailed with a polyA sequence using an enzymatic approach with *E. coli* PAP (Lucigen). The PAP reaction mixture (5–10 μL) was incubated at 37 °C for 45 min (for the preparation of RNA G1 with phosphorothioate-modified polyA), while the preparation of the RNA H1 reaction mixture was incubated for 10 min (for ATP and ATP:ATPαBH_3_ D1 49:1,24:1, and 14:1), 12 min (for ATP:ATPαBH_3_ D1 9:1), 15 min (for ATP:ATPαBH_3_ D1 5:1), 17 min (for ATP:ATPαBH_3_ D1 3:1), or 45 min (for ATP:ATPαBH_3_ D1 1:1 and ATPαBH_3_ D1 only). The reaction mixture contained: PAP buffer (50 mM Tris-HCl pH 8.0, 250 mM NaCl, and 10 mM MgCl_2_), 0.2 U/μL PAP enzyme, 10 U RiboLock RNase inhibitor, 0.5 μM RNA G or 0.2 μM RNA H, ATP, or a mixture of ATP and ATPαS D1 or ATP and ATPαBH_3_ D1 at a total concentration of 0.25 mM; the reaction was stopped by the addition of EDTA to a final concentration of 25 mM. Polyadenylated mRNA was extracted with phenol:chloroform (1:1, v/v) and precipitated with 1.25 M ammonium acetate and isopropanol at RT, followed by a washing step with 70% ethanol. Polyadenylated mRNA pellets were resuspended in 10 μL of RNase/DNase free water (Sigma). The absorbance of the mRNA sample was measured spectrophotometrically (NanoDrop 2000c), and the concentration was determined using an extinction coefficient predicted for a particular mRNA sequence by DNA Calculator (software available on www.molbiotools.com website). To assess the length of polyA tails, polyadenylated mRNAs were electrophoretically resolved in a 1% 1× TBE agarose gel between two RNA ladders (RiboRuler High Range, ThermoFisher Scientific) and analyzed using Image Lab™ Software 6.0.1 (Bio-Rad). Bands were automatically detected and the length of mRNAs was calculated using semi-logarithmic analysis with reference to the two RiboRuler RNA ladders. The polyA tail length was estimated as the difference between the length of the polyadenylated mRNA and the transcript without a polyA tail, used as a primer for PAP.

### 4.6. Purification of human CNOT7 deadenylase

The cDNA region encoding full length human CNOT7 deadenylase was subcloned into pET51b-derived expression vector with an N-terminal StrepTAGII-HA and C-terminal 10xHIS tag. Fusion protein was expressed in *E.coli* BL21(DE3) CodonPlus-RIL (Stratagene) in Super Broth Auto Induction Media (Formedium, Norfolk, UK) supplemented with ampicilin (50 μg/ml) for 48 h at 18 °C. Cells were spin down at 4000 × g at 4 °C for 15 min and stored in −20 °C. For purification, cells from 2 l of culture (60 g) were thawed, resuspended in 180 ml of lysis buffer (150 mM NaCl, 50 mM Tris pH 8.0, 10 mM imidazole, 10 mM β-mercaptoethanol, 300 mM urea, 50 mM arginine, 50 mM glutamic acid) supplemented with protease inhibitors (PMSF, pepstatin A, chymostatin, leupeptin, benzamidine) and lysosyme (0.25 mg/ml) and disrupted with EmulsiFlex-C3 high pressure homogenizer (Avestin Europe GmbH, Mannheim, Germany). Cell extract was clarified by ultracentrifugation at 140 000 x g at 4 °C for 45 min and recombinant CNOT7 protein was purified automatically using ÄKTAxpress FPLC system (GE Healthcare). The following protocol was used for automated purification: (1) nickel affinity chromatography (buffer: 150 mM NaCl, 10 mM Tris pH 8.0, 10 mM β-mercaptoethanol, 10 mM or 600 mM imidazole); (2) size-exclusion chromatography (buffer: 150 mM NaCl, 10 mM Tris pH 8.0; column: HiLoad 16/60 Superdex 200). Purity of CNOT7 deadenylase was validated on 12% PAGE. Purified protein was concentrated by ultrafiltration to 110-120 μM, flash-frozen with liquid nitrogen in storage buffer (SEC buffer supplemented with 15% glycerol and 1 mM DTT) and stored at −80 °C.

### 4.7. PolyA tail degradation assay (RNA G1 and H1)

The stability of polyA tails modified with phosphorothioate (RNA G1) or boranophosphate (RNA H1) was examined by subjecting mRNAs to deadenylation using recombinant CNOT7 deadenylase (64). A single experiment was performed as a set of simultaneous reactions for mRNAs polyadenylated by PAP and the mixture of ATP and phospho-modified analog as a substrate (e.g. ATP:ATPαS D1 marked as 9:1,4:1,3:1, 2:1, and 1:1). RNA G1 or H1 with an unmodified polyA tail (marked as 10:0) served as a positive control. Each degradation assay was performed as a single-tube reaction, containing: CNOT7 buffer (10 mM Tris-HCl pH 8.0, 50 mM KCl, 5 mM MgCl_2_, and 10 mM DTT), polyadenylated mRNA (12 ng/μL), and CNOT7 deadenylase (0.2 μg/μL). After 0, 20, 40, 60, or 120 min of incubation at 37 °C, 2.5 μL of the reaction mixture was collected and the CNOT7 deadenylase was inactivated by addition of an equal volume of loading dye (95% formamide, 50 mM EDTA, 0.025% SDS, 0.025% bromophenol blue, 0.025% xylene cyanol, and 0.025% ethidium bromide). Products of the degradation were analyzed by 1% 1× TBE agarose gel electrophoresis. Using Image Lab™ Software 6.0.1 (Bio-Rad), the degradation rate was estimated as the difference of the mRNA length before deadenylation and its length at a particular time point.

### 4.8. *In vitro* translation with rabbit reticulocyte lysate (RNA F)

The Rabbit Reticulocyte Lysate System (Promega) was used to determine the influence of internal modifications on translation efficiency of RNA F. The reaction mixture (9 μL) contained: reticulocyte lysate (4 μL), amino acid mixture without leucine (0.2 μL, 1 mM solution; the final concentration in 10 μL reaction mix was 20 μM), amino acid mixture without methionine (0.2 μL, 1 mM solution; the final concentration in 10 μL reaction mix was 20 μM), potassium acetate (1.9 μL of 1 M solution), and MgCl_2_ (0.4 μL of 25 mM solution). After 1 h of incubation at 30 °C, 1 μL of the appropriate RNA F dilution (3.0 ng/μL, 1.5 ng/μL, 0.75 ng/μL, or 0.375 ng/μL) was added and the incubation of the reaction mixture was continued at 30 °C for 1 h. The reaction was stopped by freezing in liquid nitrogen. To detect luminescence from *Firefly* luciferase, 50 μL Luciferase Assay Reagent (tricine, Mg(HCO_3_)_2_, MgSO_4_, 0.1 mM EDTA, DTT, ATP, luciferin (VivoGlo™ Luciferin, *In Vivo* Grade; Promega), and coenzyme A) was added to the sample and the luminescence was measured on a Synergy H1 (BioTek) microplate reader. The results are presented as proportions between the regression coefficients of the linear relationships between the RNA F concentration in the translation reaction (0.3 ng/μL, 0.15 ng/μL, 0.075 ng/μL, or 0.0375 ng/μL) and the corresponding luminescence signal. The relative luciferase activity was normalized to the values obtained with unmodified RNA F, calculated as mean values from three independent experiments ± standard deviation (SD).

### 4.9. Protein expression studies (RNA G, G1, H, H1, and I)

The murine immature dendritic cell line JAWS II (American Type Culture Collection (ATCC) CRL-11904) was grown in RPMI 1640 medium (Gibco) supplemented with 10% fetal bovine serum (FBS), sodium pyruvate (Gibco), 1% penicillin/streptomycin and 5 ng/mL granulocyte-macrophage colony-stimulating factor (PeproTech) at 5% CO_2_ and 37 °C. HeLa (human cervical epithelial carcinoma, ATCC CCL-2) cells were grown in DMEM (Gibco) supplemented with 10% FBS (Sigma), GlutaMAX (Gibco) and 1% penicillin/streptomycin (Gibco) at 5% CO_2_ and 37 °C. For all experiments, the passage number of studied cells was between 5 and 25. In a typical experiment, 10^4^ per well JAWS II cells were seeded into a 96-well plate on the day of the experiment in 100 μL medium without antibiotics. In the case of the HeLa cell line, 4·10 HeLa cells per well were seeded 24 h before transfection into a 96-well plate in 100 μL medium without antibiotics. Cells in each well were transfected for 16 h using a mixture of 0.3 μL Lipofectamine MessengerMAX Transfection Reagent (Invitrogen) and 25 ng mRNA in 10 μL Opti-MEM (Gibco). To assess *Gaussia* luciferase expression at multiple time points, the medium was fully removed and replaced with fresh media after 16, 40, 64 and 88 h. To detect luminescence from *Gaussia* luciferase, 50 μL 10 ng/mL h-coelenterazine (NanoLight) in PBS was added to 10 μL cell culture medium and the luminescence was measured on a Synergy H1 (BioTek) microplate reader. Total protein expression (cumulative luminescence) for each mRNA over 4 days was reported as a mean value ± SD normalized to RNA I with a template-encoded A_128_ polyA tail.

## AUTHOR CONTRIBUTIONS

D.S., M.S., J.K., and J.J. designed the study. D.S. synthesized nucleotides used in the work, obtained and analyzed RNAs A, A1, B, E, and E1, and performed all MS experiments. M.S. obtained RNAs F, F1, F2, G, G1, H, H1, and I and performed polyA tail degradation assays (RNA G1, H1). P.S. performed protein expression studies in cultured cells (RNA G, G1, H, H1, I). M.W. obtained RNA C. D.S., M.S., J.K., and J.J. wrote the first draft of the manuscript. All authors approve the final version of the manuscript.

## ACKNOWLEDGEMENTS

We thank Mariusz Czarnocki-Cieciura (International Institute of Molecular and Cell Biology, Warsaw) for providing purified recombinant human CNOT7 deadenylase.

